# TFU72 is a novel and potent DNA-PKcs inhibitor for enhancing homology-directed repair gene editing

**DOI:** 10.64898/2026.03.10.710954

**Authors:** Sridhar Selvaraj, Ludwig Schmiderer, Jacob Lamberth, Thomas Fuchss, Freja K. Ekman, Norman F. Russkamp, Matthew H. Porteus

**Affiliations:** Department of Pediatrics, Stanford University, Stanford, CA 94305, USA; Institute for Stem Cell Biology and Regenerative Medicine, Stanford University Stanford, CA 94305, USA; MilliporeSigma, St. Louis, Missouri, 2909 Laclede Ave, USA; the life science business of Merck KGaA, Darmstadt, Germany; Healthcare business of Merck KGaA, Darmstadt, Germany; Department of Genetics, Stanford University, Stanford, CA 94305, USA

**Author notes:** contributed equally (Sridhar Selvaraj, Ludwig Schmiderer).

## Abstract

Precise gene editing through homology directed repair (HDR) is one of the most versatile genome editing approaches with broad applications. Achieving high HDR gene editing efficiency is critical to realizing the full potential of this approach. Although many strategies have been explored to enhance HDR editing efficiency, inhibition of DNA-dependent protein kinase catalytic subunit (DNA-PKcs), a key component of the non-homologous end joining (NHEJ) pathway remains one of the most effective. Here we describe a novel, highly potent DNA-PKcs small molecule inhibitor, TFU72 which enhances HDR gene editing efficiency remarkably by up to 30-fold in cell lines and human primary cells. We assessed the previously reported genotoxic outcomes associated with DNA-PKcs inhibition such as off-target mutations, chromosomal translocations and large deletions and describe approaches to mitigate these outcomes to safely enhance HDR gene editing efficiency with TFU72. This optimized approach enables broad application of TFU72 for HDR-based precise gene editing applications in both therapeutic and research settings.

## Introduction

Precise genome modification is an essential technology in the development of new therapies for genetic diseases, the development of new modalities to treat common diseases, and in the search for new biological insights. The ability to accurately correct disease-causing mutations has the potential to transform the treatment of monogenic disorders^1,2^. One of the most versatile tools for installing genetic modifications is CRISPR-Cas9 in combination with a repair template. The two-component Cas9-gRNA ribonucleoprotein is a programmable, efficient, and site-specific nuclease^3–5^. The gRNA component guides Cas9 to its cognate site in the targeted DNA where it induces a double-strand break (DSB) by DNA cleavage^6^. Precise editing can be accomplished by providing an exogenous donor template to make the intended modifications in the genome. However, the efficiency of these precise editing pathways is often limited by the competition between multiple repair pathways for fixing the DSBs^7^. Among the multiple pathways that compete to fix the break, only homology-directed repair (HDR) and certain other donor-dependent pathways, can integrate the provided repair template correctly^8^. The other pathways can result in precise rejoining of the cut DNA that restores the original sequence, or in deletions and insertions (INDELs) that can impair gene function^9^.

Within seconds^10,11^ of DSB formation, two molecular machines compete for capture of the broken DNA ends: the Ku70/80 complex, which leads to error-prone non-homologous end joining (NHEJ), and the MRN complex, which initiates the DNA resection required for accurate HDR^12^. The NHEJ pathway which is active in all cell cycle phases is initiated by the highly abundant Ku70/80 which forms a ring around the DNA ends and recruits DNA-PKcs to generate the stable DNA-PK holoenzyme^13–15^, which coordinates end processing and recruitment of XRCC4–Ligase IV that seals the break^16^. NHEJ pathway often repairs the DSB error-free which leads to further rounds of Cas9-induced DSBs and repair during genome editing. However, after multiple rounds, the NHEJ pathway leads to errors in the form of INDELs^17^. In contrast, HDR requires several hours and operates primarily during the S and G2 phases of the cell cycle^17–21^. For successful completion, extensive DNA processing which includes end resection, RPA coating, and RAD51 filament formation are required^22,23^. Even when cells attempt HDR, a third, DNA polymerase theta-dependent pathway called microhomology-mediated end joining (MMEJ), competes for resected DNA ends and generates INDELs if successful^24^. During genome editing, the HDR pathway has been shown to outcompete the INDEL outcomes generated by MMEJ when the appropriate exogenous HDR donor template is delivered to the cells. However, the NHEJ generated INDEL outcomes are not as efficiently outcompeted by precise HDR editing. Thus, NHEJ INDELs are usually the primary outcome of DNA repair in genome editing experiments^25,26^.

DNA-PKcs is a druggable bottleneck for NHEJ. When it is inhibited, Ku proteins may still bind the DNA ends but cannot complete the repair^27,28^. This opens a time window for the HDR machinery to engage^29^. Our previous work identified the DNA-PKcs inhibitor AZD7648 as a potent pathway switching agent that can improve HDR rates dramatically, enhancing template integration from 30% to over 90% at certain genomic sites^30^. Combined inhibition of DNA-PKcs and DNA polymerase theta can further improve the efficiency^31^. Genome editing with AZD7648 has been shown to have increased off-target editing^30^, on-target large deletions and translocations between on and off-target sites^32^. This necessitates the need to explore novel DNA-PKcs inhibitors and assess whether they could potentially have less unintended genotoxic outcomes than AZD7648.

Here we report TFU72, a novel small molecule DNA-PKcs inhibitor which has a lower IC50 value than AZD7648 (53 nM vs. 180 nM). Kinome profiling revealed that TFU72 has a distinct off-target kinase-inhibition profile and spares several kinases that are mildly affected by AZD7648 inhibition. TFU72 has comparable HDR-enhancing ability in multiple primary human cell types that are relevant for ex vivo gene therapy: CD34+ hematopoietic stem and progenitor cells (HSPCs), T cells, and induced pluripotent stem cells (iPSCs). We find that for gene editing with TFU72, reducing the compound incubation time and the use of HiFi Cas9 over WT Cas9 largely mitigates the off-target editing, large on-target deletions and translocation events. Thus, TFU72 is a valuable addition to the genome editing toolbox for precise HDR editing in cell lines and primary cells for research and therapeutic applications.

## Results

### Characterization of the specificity profile of TFU72

DNA-PKcs belongs to the family of Phosphoinositide 3-kinases (PI3K), and it shares structural homology with other members of the family^33^. Previously reported small molecule inhibitors of DNA-PKcs have all been shown to exhibit varying degrees of cross-reactivity to other kinases in the PI3K family^34^. To characterize the specificity of the novel DNA-PKcs inhibitor TFU72, we assessed the biochemical inhibition of the various kinases in the PI3K family. We found that the cellular half maximal inhibitory concentration (IC_50_) against DNA-PKcs was 53 nM for TFU72 while biochemically it had an IC_50_ of around 300 nM for PI3Kα, PI3Kβ, PI3Kδ and 1940 nM for PI3Kγ. For the other kinases we assessed such as ATM, ATR and mTOR, we found minimal cross reactivity with a biochemical IC_50_ of more than 10,000 nM. In comparison, AZD7648 had a cellular IC_50_ of 180 nM for DNA-PKcs, showed partial inhibition of PI3Kγ (IC_50_: 300 nM) and high inhibition of ATM (IC_50_: 83 nM) (Fig. 1A). To further confirm the specificity of TFU72, we tested the compound against a panel of 513 human kinases which includes wildtype (WT) and mutant enzymes (methods description included in Supplemental information). We used the compound at a concentration of 1 μM which we found to be the optimal concentration for HDR editing as shown below. The top hit in the kinase panel for TFU72 was DNA-PKcs with more than 98% inhibition relative to a previously characterized positive control compound. We found that the mutant and WT forms of PI3Kα were the other kinases which were inhibited by TFU72 at more than 50% (Fig. 1B). Several other kinases were partially inhibited at 20-50% but most of them were inhibited at less than 30% relative to a positive control (Fig. S1A). In comparison, AZD7648 also showed the top hit to be DNA-PKcs while it inhibited PI3Kα, PI3Kγ and ATM at above 50% (Fig. 1C). mTOR and PI3Kα variants were some of the kinases inhibited at less than 50% by AZD7648 (Fig. S1B). The full kinase panel testing data is included in Table S2. Thus, TFU72 showed efficient DNA-PKcs inhibition and very minimal inhibition of ATM and ATR kinases which play a key role in homology directed repair (HDR)^35^ making it a potentially good candidate for HDR enhancing applications.

**Figure 1.**
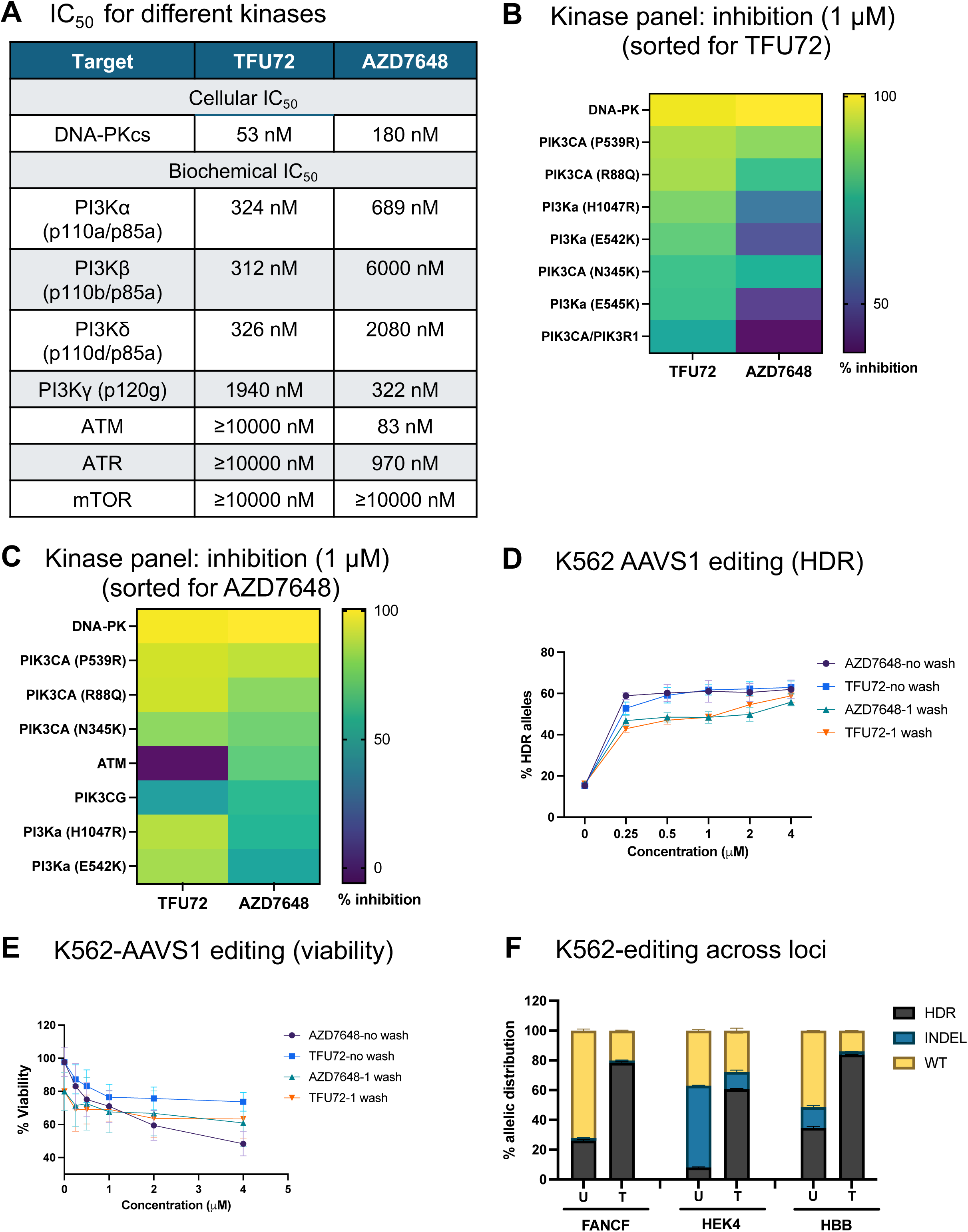
Specificity profile and HDR enhancing ability of TFU72 in comparison with AZD7648. **A.** Specificity profile of TFU72 and AZD7648 against kinases of the PI3K and PIKK families. Data is shown as half maximal inhibitory concentration (IC_50_) against (cellular IC_50_ for DNA-PKcs, biochemical IC_50_ for other kinases) different kinases. **B-C.** Heatmap showing the top kinases inhibited by TFU72 and AZD7648 (both at 1 µM) following testing on a human kinase panel. The data is shown in descending order sorted for kinases inhibited at 50% or above relative to a target-specific inhibitor for TFU72 (**B**) or AZD7648 (**C**). **D-E.** Allelic HDR editing frequencies at AAVS1 locus determined by NGS analysis at D5 post RNP and ssODN editing with different concentrations of TFU72/AZD7648 as indicated (**D**). The compounds were removed either at 24 hours post editing (1 wash) or left in the cells for 5 days (0 wash). Viability of the gene edited cells as measured by CellTiter-Glo® 2.0 Assay is shown as percentage relative to the untreated sample (gene editing without compound) (**E**) (n=2). **F.** Allelic distribution of HDR, INDEL and WT frequencies in K562 cells edited across different gene loci (FANCF, HEK4, HBB) with or without TFU72 treatment based on NGS analysis. U denotes untreated (editing without compound), T denotes editing with TFU72 (1 µM) (n=2).

### TFU72 enhances HDR editing in immortalized cell lines

We have shown previously that AZD7648 was the most potent and selective compound for enhancing HDR gene editing efficiency^30^. Thus, we tested the HDR enhancing potential of TFU72 in comparison with AZD7648 throughout this study. First, we compared the compounds for HDR gene editing with Cas9/sgRNA ribonucleoprotein (RNP) and single-stranded oligodeoxynucleotide (ssODN) HDR donor in immortalized cell lines. To identify the optimal concentration, we tested different doses of the compounds (0.25, 0.5, 1, 2 and 4 μM) for HDR gene editing at AAVS1 locus in K562 cells with and without washing off the compounds at 24h post nucleofection. We found the minimum optimal concentration to be between 0.25 to 1 μM for both compounds and observed that the HDR efficiency was higher by 1.1- to 1.3-fold in cells incubated with the compound either with or without the 24h wash step (no wash) (Fig. 1D). Correspondingly, we observed a decrease in the allelic frequency of WT (∼4- to 8-fold) and INDELs (∼1.5- to 2-fold) in the cells edited with the compounds (Fig. S2A). Viability assessment showed that cells edited with TFU72 had improved viability relative to that of AZD7648 in the no wash condition, but the viability was similar in the 1 wash condition (Fig. 1E). We performed the dose titration at a second gene locus, BCL11A in K562 cells and further confirmed that the minimal optimal concentration was between 0.25 to 1 μM for both the compounds (Fig. S2B). At the chosen concentration of 1 μM, we tested the TFU72 for editing three other gene loci and observed a HDR improvement of ∼2- to 3-fold at FANCF and HBB loci, more than 10-fold at HEK4 (a previously described non-coding genomic target site) (Fig. 1F). Next, we assessed whether TFU72 can enhance HDR editing in other immortalized cell lines, HEK293 and U2OS. In HEK293, we observed a dramatic improvement in HDR at both loci tested (1.6% to 28% at HEK4, 1.8% to 48% at HBB). In U2OS cells, we observed no detectable HDR without treatment at HEK4 but TFU72 treatment yielded ∼6% HDR (Fig. S2C and D). Thus, TFU72 enhances HDR gene editing efficiency across different loci in various immortalized cell lines.

### TFU72 enhances HDR editing in human primary cells

To assess whether TFU72 can enhance HDR in therapeutically relevant cell types, we tested it for gene editing in human iPSCs, HSPCs and T cells. We used the previously optimized and clinically relevant gene editing platform of HiFi Cas9 RNP^36^ and AAV6-based HDR donor template delivery for all our gene editing studies in human primary cells^37,38^. First, we performed a dose titration study comparing TFU72 and AZD7648 for HDR gene editing to knock-in a 6 bp sequence at the CCR5 locus (two stop codons) (Fig. S3A). In all three cell types, we observed that both compounds enhanced HDR efficiency to a similar extent at the optimal doses of 0.5 to 1 μM. HDR efficiency improved by more than 3-fold in iPSCs with the compound treatment (mean of 28% to 97%) and the minimum optimal concentration was found to be 0.5 μM for AZD7648 and 1 μM for TFU72 (Fig. 2A). Since the gRNA we used for these studies generates predominantly NHEJ INDELs over HDR, we found that the compound treatment efficiently reduced the NHEJ INDEL frequency to improve the HDR (Fig. 2B). Using an MTT cell viability assay, we observed that the viability of iPSCs edited with TFU72 was slightly higher than those edited with AZD7648, but this difference was not statistically significant (Fig. S3B). In HSPCs, both compounds enhanced HDR efficiency by 4-fold and the minimum optimal concentration was 0.5 μM (Fig. 2C). Correspondingly, we observed a reduction in the allelic NHEJ frequency in the cells edited with the compounds (Fig. 2D). Upon measuring the viable cell count, we noticed that both compounds had minimal effect on the viability especially at concentrations less than 2 μM (Fig. S3C). To further confirm the effect of the compounds on viability and evaluate the differentiation potential of the CCR5 gene edited HSPCs (Fig. S3D), we performed a colony forming units (CFU) assay. We detected a decrease in the total number of colonies in the HDR edited cells without the compounds when compared to the Mock samples as described previously^39,40^ (Fig. S3E). This drop in colony number was maintained in the cells edited with AZD7648 and we observed a slight increase in the number of colonies in the TFU72 sample, but the difference was not statistically significant. The differentiation potential was not altered in the gene edited cells with or without the compounds as indicated by the similar colony type distribution (granulocyte, erythroid, macrophage, megakaryocyte (GEMM), granulocyte and monocyte (GM), and burst forming unit erythroid (BFU-E)) (Fig. S3E). In T cells, we found the minimum optimal concentration for the two compounds to be 1 μM with a marked increase in allelic HDR efficiency from ∼1% to ∼80% (Fig 2E). At the lower concentrations, we detected that the TFU72 outperformed AZD7648 in improving the HDR efficiency. Like in iPSCs and HSPCs, we observed a corresponding decrease in the allelic NHEJ frequency with the compound treatment. Additionally, we also noticed there was a small frequency of alleles with MMEJ INDELs only in the cells edited with the compounds and this outcome was unique to the T cells (Fig. 2F).

**Figure 2.**
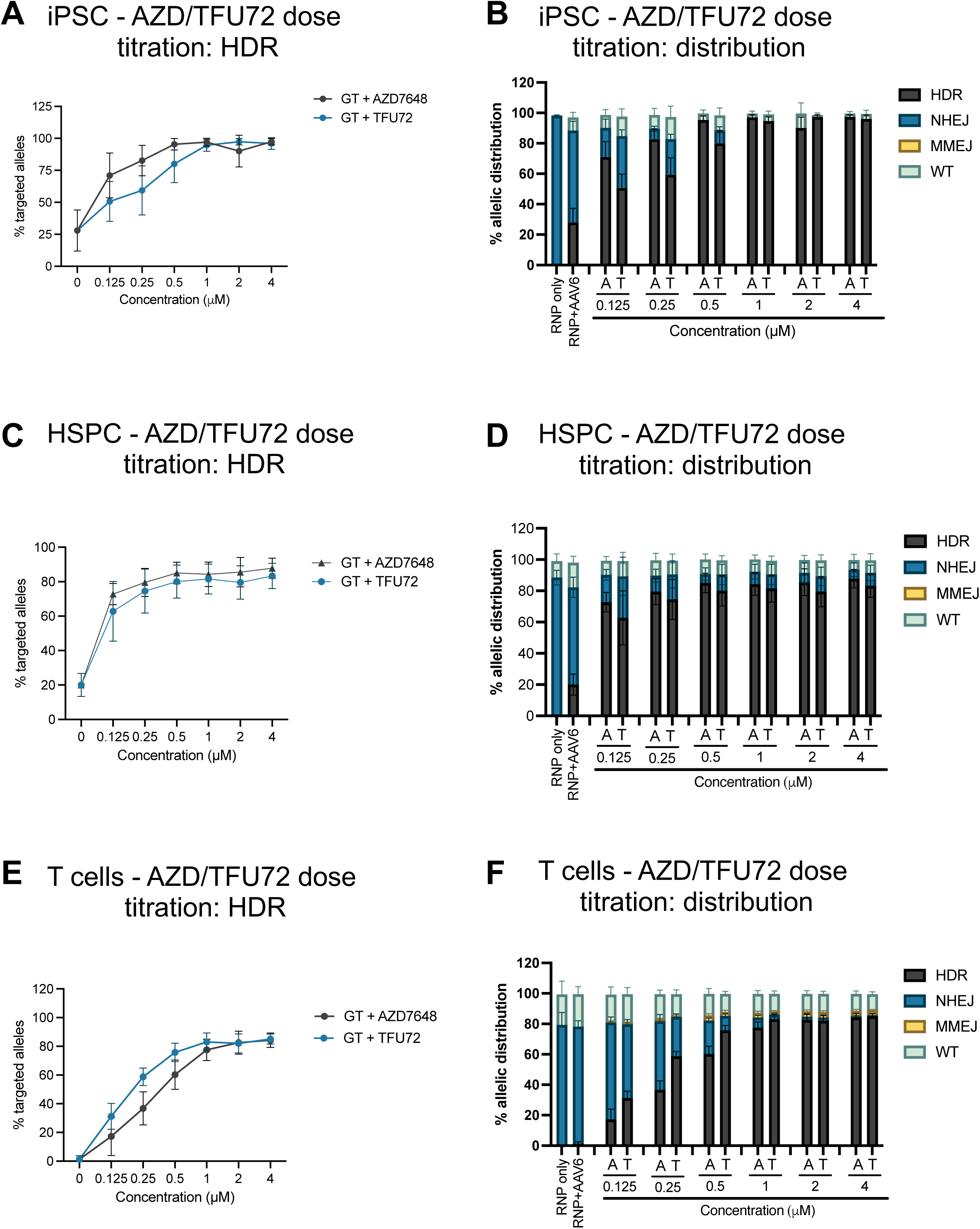
Dose titration of TFU72 for enhancing HDR editing in iPSC, HSPC and T cells. iPSC, HSPC and T cells were edited at CCR5 locus using RNP and AAV6 gene editing for knock-in of short sequence (two stop codons) with different concentrations of AZD7648 and TFU72 as indicated. HDR, NHEJ, MMEJ and WT allelic frequencies were calculated at D3 post editing using Sanger Sequencing and ICE analysis (n=3). **A, C, E**. Scatter plots show the allelic HDR frequency in iPSC (**A**), HSPC (**C**) and T cells (**E**). **B, D, F**. Bar graphs show the distribution of allelic HDR, NHEJ, MMEJ and WT frequencies in iPSC (**B**), HSPC (**D**) and T cells (**F**). GT denotes gene targeting (HDR) with RNP, AAV6 editing. A and T denote AZD7648 and TFU72, respectively.

Next, we tested TFU72 for HDR editing individually across three different gene loci (CCR5, HBB and STING1) with two gRNAs each in iPSCs, HSPCs and T cells. For this, we tested one predominantly NHEJ guide RNA (R11-CCR5, sg3-HBB) and one MMEJ guide RNA (sg1-CCR5 and R02-HBB) each at CCR5 and HBB loci. At the STING1 locus, we chose one predominantly NHEJ guide RNA (sg3) and one low INDEL frequency guide RNA (sg5). For these studies, we used AAV6 HDR templates for knock-in of short sequences. AAV6 template for CCR5 locus was designed to knock-in two stop codons as described above (Fig. S3A). For the HBB locus, we used the AAV6 HDR template for editing sickle cell disease (SCD) mutation and the template also introduces silent mutation at the gRNA target site (Fig. S4A). The STING1 AAV6 HDR template was designed to knock-in a point mutation (V155M) along with the silent mutations at the gRNA target site (Fig. S4B). In iPSCs, we tested TFU72 and AZD7648 at two concentrations each (0.5 and 1 μM) for editing at the three loci and we used the compounds at the optimal concentrations of 0.5 μM and 1 μM each in HSPCs and T cells, respectively. At the CCR5 locus, we observed that the editing with the NHEJ guide RNA (R11) recapitulated the HDR enhancement with reduction in NHEJ INDEL frequency observed in the dose-titration studies across all three cell types (Fig. 3A, B, C, Fig. S4C, D, E). With the second guide RNA (sg1) at CCR5 locus, we detected high HDR efficiency in the RNP/AAV6 samples but the treatment with the compounds improved the HDR further in the iPSCs (70% to >95%) (Fig. 3A). The RNP only sample for the CCR5-sg1 gRNA in iPSCs showed high MMEJ INDEL frequency which has been shown previously to correlate with high HDR frequency upon editing with the donor template^29^ (Fig. S4C). Interestingly, we observed that the CCR5-sg1 gRNA produced no INDELs in the RNP-only edited samples and low HDR frequency in the RNP/AAV6-edited samples in HSPCs and T cells. However, we found that the compounds dramatically improved the HDR frequency in HSPCs and T cells (2% to 30% in HSPCs and 0.33% to 48% in T cells) (Fig. 3B, C). Editing with the NHEJ gRNAs at the HBB and STING1 loci (sg3 for both) resulted in significant improvement in HDR efficiency with the compounds in all three cell types (Fig. 3A, B, C) and correspondingly we noticed a drastic reduction in the NHEJ INDEL frequency (Fig. S4C, D, E). Upon editing with the MMEJ gRNA at HBB locus (R02), we observed high HDR efficiency without treatment, but the compounds did enhance the HDR further in all three cell types (76% to >90% in iPSC, 66% to >80% in HSPC and 24% to >50% in T cells) (Fig. 3A, B, C). The low to no INDEL frequency guide RNA (sg5) editing at STING1 locus yielded low HDR but the compound treatment dramatically improved the HDR efficiency in all three cell types (Fig. 3A, B, C). This data recapitulated the observation in our previous study on editing with the same sg5 STING1 gRNA and AZD7648 indicating that TFU72 can also enhance HDR editing efficiency dramatically even with a seemingly inactive low INDEL frequency gRNA^30^. This observation indicates that genome editing with the seemingly inactive gRNAs does induce double strand breaks which are repaired error-free by NHEJ and blocking this repair with the DNA-PKcs inhibition allows for precise HDR editing with the exogenous donor template^30^. Overall, we found that TFU72 enhances HDR efficiency across genomic loci and gRNAs at similar efficiency as AZD7648.

**Figure 3.**
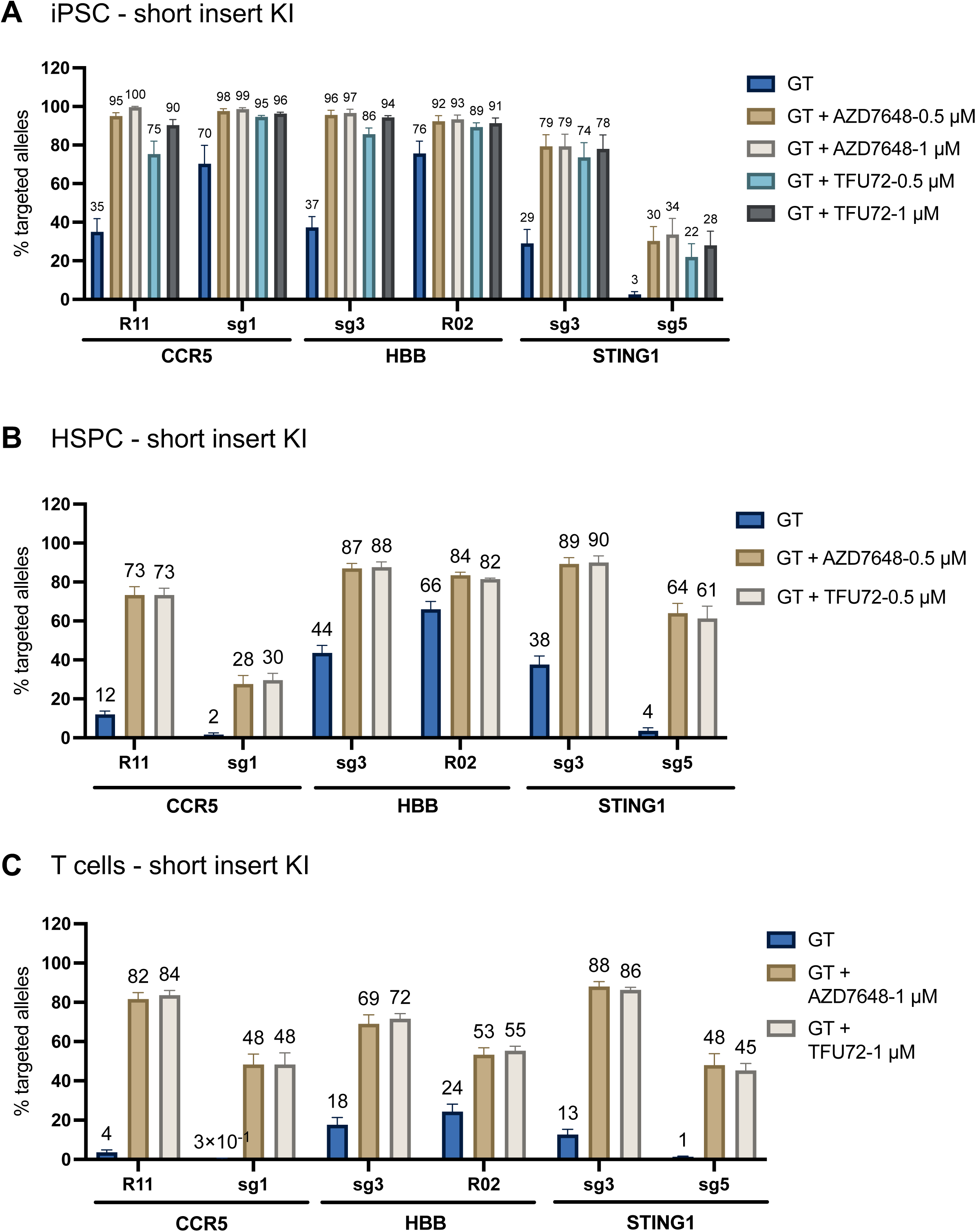
HDR editing with TFU72 across different loci and gRNAs in iPSC, HSPC and T cells. A-C. iPSC (**A**), HSPC (**B**) and T cells (**C**) were edited individually at CCR5, HBB and STING1 loci with two different gRNAs at each loci using RNP and AAV6 gene editing for the knock-in of short sequence with AZD7648 and TFU72 at the indicated concentrations. Bar graphs show allelic HDR frequencies (% targeted alleles) measured at D3 post editing using Sanger Sequencing and ICE analysis (n=3). GT denotes gene targeting (HDR) with RNP, AAV6 editing. AAV6 HDR template for CCR5 locus was designed to knock-in two stop codons at the target site. At the HBB locus, the AAV6 HDR template was designed to knock-in silent mutations and correction of the sickle cell disease mutation (E6V). For the STING1 locus, the AAV6 HDR template was designed to knock-in a point mutation (V155M) along with silent mutations at the gRNA target site.

### TFU72 enhances large insert knock-in HDR

The RNP/AAV6 gene editing platform is typically used for the integration of large multi-kb sequences. So, we assessed whether TFU72 enhances HDR for the knock-in of large inserts (1kb or larger in length) independent of the amount of AAV6 donor template used by titrating the AAV multiplicity of infection (MOI: 500, 1000, 2500 and 5000). We used the NHEJ guide RNAs to edit individually at CCR5 (R11), HBB (sg3) and STING1 (sg3) loci in iPSC, HSPC and T cells. The donor templates for CCR5 and HBB loci were designed to contain a knock-in sequence of the Ubiquitin C promoter, GFP cDNA with the bGH polyA signal sequence (∼2.2 kb) (Fig. S5A, B) while the insert sequence for the STING1 locus contained the PGK promoter, GFP cDNA and the sv40 polyA signal sequence (∼1.3 kb) (Fig. S5C). In iPSCs, TFU72 enhanced HDR editing at similar levels as AZD7648 at all three loci both in terms of allelic frequency (% alleles with HDR) and cellular frequency (% cells positive for HDR (GFP)) measured by ddPCR and flow cytometry, respectively (Fig. 4A, Fig. S6A). Importantly, at the lowest MOI tested (500), the HDR efficiency was higher with treatment when compared to the HDR without treatment at the highest MOI (5000) indicating that the HDR improvement with TFU72 is efficient even at low MOIs. We did observe a positive correlation of HDR frequency with the MOI in iPSCs (Fig. 4A, Fig. S6A). Correspondingly, NHEJ INDEL frequency decreased in TFU72-treated iPSCs, while the frequency of WT alleles increased and a small fraction of MMEJ INDEL alleles appeared that were not detected in untreated cells (Fig. S6B). In HSPCs, the allelic and cellular HDR editing frequencies with the compound treatment improved at all three loci with a positive correlation to the MOI. We detected a fold increase of ∼2 to 6 with the treatment for allelic HDR at the lowest MOI (500) and ∼1.6- to 4-fold increase at the highest MOI (5000) (Fig. 4B). The cellular HDR frequency improved by 1.6- to 6-fold at the lowest MOI (500) and by 1.6- to 3.6-fold at the highest MOI (5000) with the treatment (Fig. S6C). Without the treatment, we noticed very low to no MMEJ INDELs at the lower MOIs (500 and 1000), but the treatment led to an increase in the frequency of MMEJ INDELs by more than 6-fold in some cases. At the higher MOIs, this MMEJ INDEL difference was minimal as the HDR frequencies improved further (Fig. S6D). In T cells, we observed a substantial improvement in the allelic and cellular HDR frequencies with the compound treatment by up to 7- and 6-fold, respectively (Fig. 4C, Fig. S6E). The compound treatment also led to a significant increase in the MMEJ INDEL frequency corresponding to the improvement in HDR and decrease in the NHEJ INDEL frequency in T cells (Fig. S6F). In summary, TFU72 enhances the HDR frequency for large sequence knock-in with different AAV6 MOIs at similar efficiency as AZD7648 across different genomic loci in iPSCs, HSPCs and T cells.

**Figure 4.**
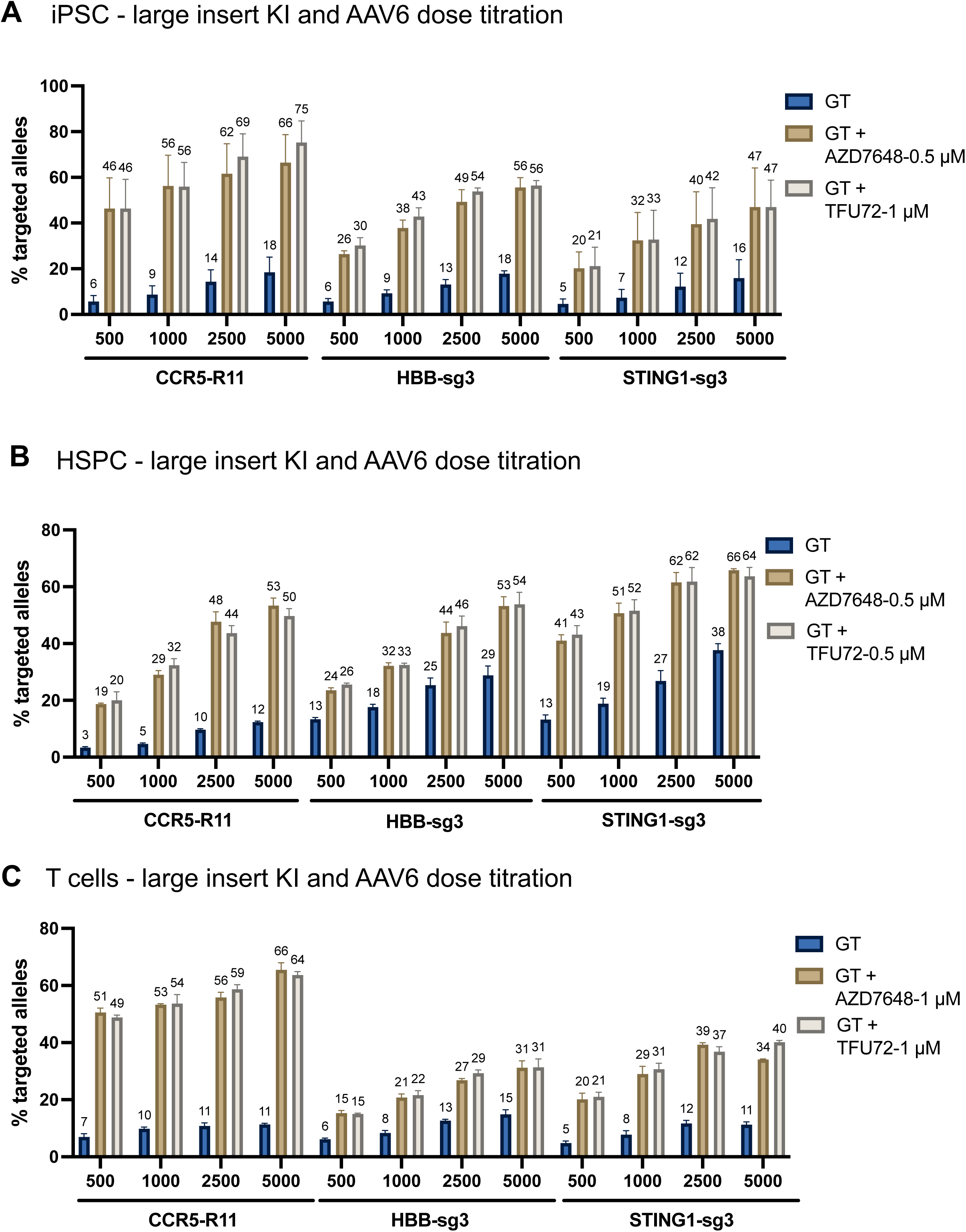
HDR editing with TFU72 for large insert KI with AAV6 dose titration in iPSC, HSPC and T cells. **A-C**. iPSC (**A**), HSPC (**B**) and T cells (**C**) were edited at CCR5, HBB and STING1 loci individually using RNP and AAV6 gene editing for the knock-in of multi-kb sequence with AZD7648 and TFU72 at the indicated concentrations and at different AAV6 doses, multiplicity of infection (MOI): 500, 1000, 2500, 5000. Bar graphs show allelic HDR frequencies (% targeted alleles) measured at D3 post editing using ddPCR analysis (n=3). GT denotes gene targeting (HDR) with RNP, AAV6 editing. AAV6 HDR templates for CCR5 and HBB loci were designed to knock-in a 2.2-kb sequence consisting of UBC promoter driven GFP followed by a bGH polyA signal sequence. For the STING1 locus, the AAV6 HDR template was designed to knock-in a 1.4-kb sequence consisting of PGK promoter driven GFP followed by a sv40 polyA signal sequence.

### Potential genotoxic outcomes and mitigation strategies

Gene editing with DNA-PKcs inhibition has been previously described to increase the frequency of unintended genotoxic outcomes such as off-target mutations, translocations and large deletions^30,32^. We assessed these outcomes in HSPCs edited at HBB locus with the SCD donor template described above. For these studies, we performed a comparison between HSPCs edited with HiFi and WT Cas9 proteins as some of the previous studies used the WT Cas9. We detected similar levels of HDR editing in HiFi and WT Cas9 edited samples (Fig. S7A). Off-target INDELs were assessed in HSPCs edited with or without the compounds at the previously characterized off-target 1 (OT1) site in Chromosome 9^41^. We found that treatment with TFU72 and AZD7648 led to a 2 to 3-fold increase in the off-target INDELs in cells edited with HiFi and WT Cas9 proteins. This data correlated with the results reported in our previous study using AZD7648^30^. However, we observed that editing with HiFi-Cas9 with or without treatment resulted in more than 10-fold lower off-target INDELs than the corresponding WT-Cas9 samples as reported previously^42^ (Fig. 5A). Correspondingly, we also observed more than 5-fold increase in the translocation events between the on-target and the OT1 off-target sites in WT-Cas9 edited samples relative to the HiFi Cas9 as measured by ddPCR. The compound treatment did increase the frequency of translocation events by 3- to 4-fold relative to the corresponding untreated samples (Fig. 5B). To characterize the large deletions, we used large amplicon sequencing by Oxford Nanopore sequencing. In HiFi Cas9-edited cells, we observed that the RNP-only editing resulted in higher frequency of large deletions than the RNP/AAV6 HDR-edited cells indicating that HDR editing mitigates the large on-target deletions. Upon treatment with the compounds, we detected a 2- to 3-fold increase in the frequency of large deletions of the sizes 1 kb to 3 kb, 3 kb to 5 kb and above 5 kb (Fig. 5C). We detected a similar pattern of large deletions in WT-Cas9 edited cells, but we only detected an increase in the frequency of large deletions of the sizes 1 kb to 3 kb in the treated cells (Fig. S7B).

**Figure 5.**
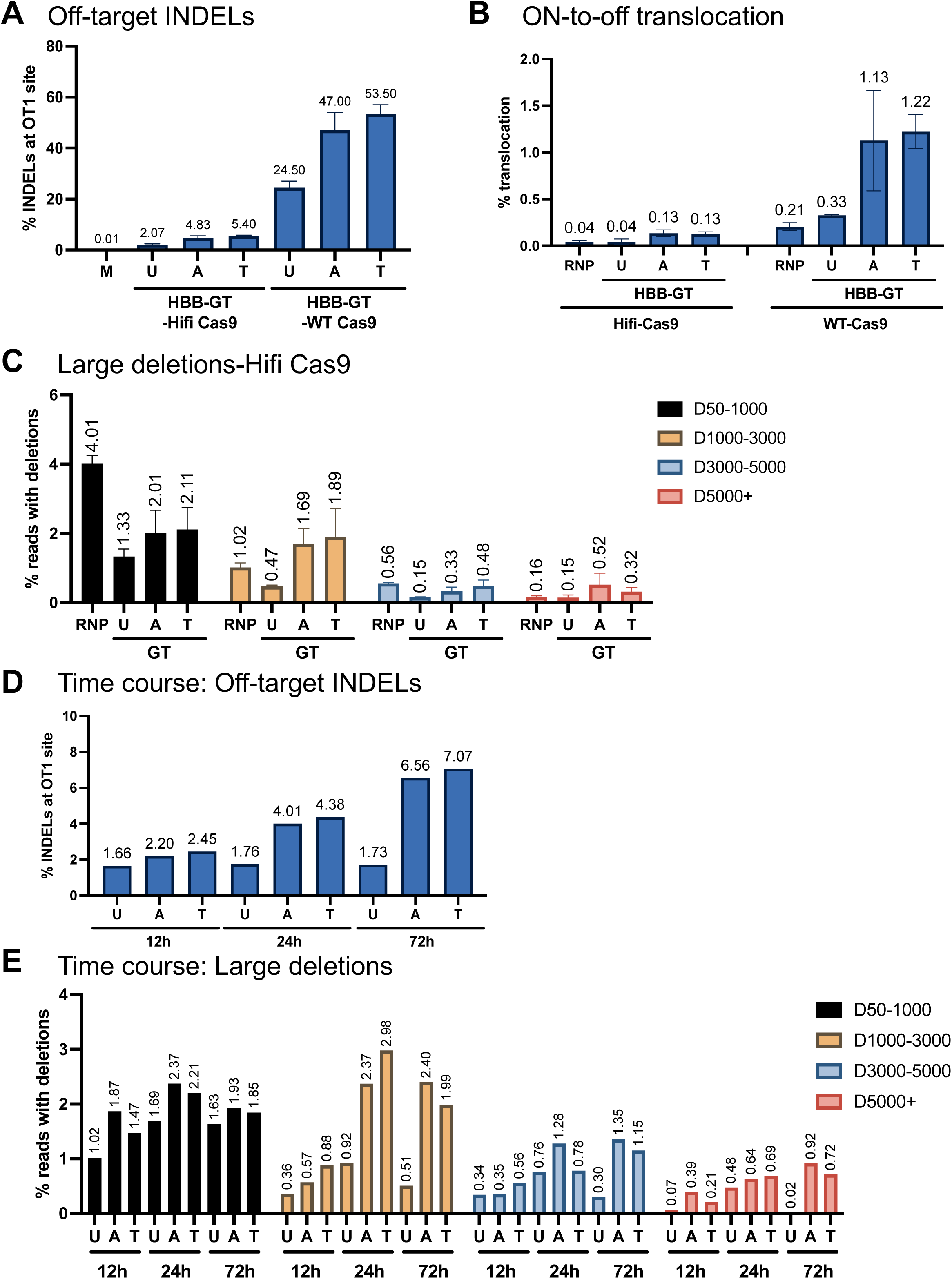
Assessment of potential genotoxic outcomes post HDR editing with TFU72 in HSPCs. HSPCs were edited at HBB locus using RNP, AAV6 (Hifi or WT Cas9) with AZD7648 (A, 0.5 µM) or TFU72 (T, 0.5 µM) or no treatment (U). Mock electroporated cells (M) were used as a negative control. GT denotes gene targeting (HDR) with RNP, AAV6. **A**. At D3 post editing, a previously characterized off-target site at Chr9 (OT1) was PCR amplified from genomic DNA and sequenced by NGS. Bar graphs show INDEL frequency at the OT1 site as determined by the CRISPResso2 tool analysis of the NGS data (n=2). WT Cas9 edited samples were assessed for the INDELs using Sanger Sequencing and ICE analysis. **B**. Translocation between on-target (HBB) site and OT1 off-target site was assessed by ddPCR analysis and data is shown as percentage of translocation which is the sum of 4 different translocation outcomes (n=2). **C**. HSPCs edited at HBB loci with Hifi Cas9 were assessed for the frequency of large deletions at the on-target site using Nanopore sequencing of a 10-kb PCR amplicon. Bar graph shows the frequency of reads with deletions of the sizes 50-1000, 1000-3000, 3000-5000 and above 5000 bp (n=3). **D-E.** HSPCs edited at HBB locus using RNP, AAV6 with different incubation times of AAV6 (U), AZD7648+AAV6 (A) and TFU72+AAV6 (T) were assessed for the off-target INDELs at the OT1 site as described above. **D.** Bar graphs show the INDEL frequency at the OT1 site at D3 post editing in samples with 12h, 24h and 72h incubation times (n=1). **E.** Bar graphs show the frequency of reads with deletions of the sizes 50-1000, 1000-3000, 3000-5000 and above 5000 bp as determined by Nanopore sequencing described above (n=1).

Next, we tested whether the potential genotoxic outcomes could be reduced by altering the TFU72 incubation time of 0 to 24h post transfection either by delaying the start time or by reducing the incubation time. The rationale for delaying the start time is based on previous studies describing that the INDEL events occur starting at around 4 to 8h post transfection^29^. We tested different start times for the incubation such as 2 to 24h, 4 to 24h, 6 to 24h and 8 to 24h post transfection. At these modified start times, we observed that the improvement in HDR efficiency with TFU72 is largely maintained in 2 to 24h and 4 to 24h samples while the 6 to 24h and 8 to 24h samples showed a smaller improvement (Fig. S7C). However, delaying the treatment start time to 4h or 8h post gene editing did not mitigate the increase in the off-target INDELs (Fig. S7D). As an alternative, we tested whether reducing the compound incubation times to 4h, 8h and 12h (start time of 0h) post transfection would mitigate the off-target INDELs and large deletions. We also included a 72h incubation time to determine whether there is an incubation time dependence for these outcomes. Since reducing the compound incubation times by earlier media change would also lead to reduced AAV6 incubation times, we compared the HDR editing to the untreated samples with the corresponding AAV6 incubation times. In the untreated samples, we observed that the HDR frequency showed a positive correlation with the AAV6 incubation times and the 72h incubation showed the highest HDR frequency. Upon compound treatment, the HDR frequency improved in all the incubation times relative to the corresponding untreated samples (Fig. S7E, F). We chose to assess the off-target INDELs and large deletions in the 12h, 24h and 72h samples as the lower incubation times showed minimal HDR improvement (Fig. S7F). Off-target INDEL frequency showed a compound incubation time dependent increase with the 72h samples showing more than a 3.5-fold increase. Importantly, the 12h compound incubation largely mitigated the increase in the off-target INDEL frequency when compared to our standard 24h incubation (Fig. 5D). Correspondingly, we detected that reducing the compound incubation time to 12h also decreases the frequency of large deletions when compared to the 24h and 72h incubation (Fig. 5E). Thus, TFU72 treatment led to an increase in the frequency of off-target INDELs, translocation and large deletions at similar levels as AZD7648. But the extent of these outcomes was largely mitigated upon editing with HiFi-Cas9 as opposed to the WT-Cas9. Reducing the compound incubation times to 12h post transfection further mitigated the off-target INDELs and large deletions.

Overall, these results indicate that TFU72 is a potent DNA-PKcs inhibitor which enhances HDR gene editing efficiency remarkably in human cell lines and primary cells. Using an optimized methodology, the genotoxic outcomes associated this approach can be largely mitigated.

## Discussion

Despite the development of novel genome editing technologies for enabling targeted genomic integration, HDR remains the most versatile and precise method for scarless integration of short to large sequences^7^. However, the efficiency of HDR is often low as DSBs are preferentially repaired by the NHEJ pathway which leads to INDELs^29^. The efficiency of HDR has been significantly improved by the application of the optimal delivery platform of Cas9 RNP and AAV6-based HDR donor template delivery^37,38^. However, the allelic HDR efficiency remains quite variable between 20 to 60% across loci and guide RNAs typically due to high NHEJ INDEL outcomes^38^. Several targets have been probed for the inhibition of NHEJ pathway to enhance the efficiency of HDR gene editing^7^. Among these, DNA-PKcs has been shown to be one of the most reliable HDR-enhancing targets following small molecule-based transient inhibition during gene editing. Many small molecule inhibitors have been developed for DNA-PKcs inhibition with varying efficacy in the recent years for anti-cancer therapeutics^43^. In a previous study, we found AZD7648 to be the most potent compound for enhancing HDR gene editing efficiency among the commercially available compounds^30^. Here we describe a novel DNA-PKcs inhibitor, TFU72 which showed a more favorable specificity profile than AZD7648 and similar HDR-enhancing activity as AZD7648. TFU72 had much lower cross-reactivity to ATM and ATR kinases which are key proteins in the HDR pathway based on biochemical and cellular assays. Importantly, we observed that TFU72 could be potentially less toxic than AZD7648 for gene editing. TFU72 improved HDR robustly in different immortalized cell lines upon editing with Cas9 RNP and ssODN HDR template and it was also feasible to culture cells without washing off the compound for up to 5 days post transfection with no differences in cell viability.

In therapeutically relevant human primary cells such as iPSC, HSPC and T cells, we observed a consistent improvement in HDR efficiency with a 24h treatment of TFU72 post gene editing with the RNP/AAV6 platform across loci and gRNAs. While using seemingly inactive gRNAs which produce low to no INDELs, we detected an increase in HDR efficiency with TFU72 by more than 30-fold which was comparable with that of AZD7648. This is particularly important as gene editing with these gRNAs may result in low off-target mutations and other potential genotoxic outcomes. TFU72 enhanced HDR efficiency for integration of sequences ranging in length from short to multi-kb in human primary cells. HDR-enhancing ability of TFU72 was maintained even while editing with lower doses of AAV6 HDR template indicating that the cells need not be saturated with the template to achieve high HDR. Gene editing with lower doses of AAV6 has been shown to be associated with reduced toxicity and improved cell health due to reduction in the p53 signaling in HSPCs^21,44^.

HDR-enhancing ability of AZD7648 has been shown to be associated with increased potential genotoxic outcomes such as off-target INDELs, translocations and large on-target deletions raising concerns about the safety of this approach^30,32^. These outcomes can be overcome in iPSCs as they occur at very low frequencies, making it feasible to select HDR edited clones without these genotoxic outcomes following single cell cloning. However, it is important to address these outcomes in primary cells which are used as a bulk gene edited population for downstream research and therapeutic applications. Upon assessing these outcomes in HSPCs, we found that TFU72 had a similar effect in increasing off-target INDELs, translocations and large on-target deletions as AZD7648 indicating that inhibition of DNA-PKcs and thereby the NHEJ pathway is the underlying cause. However, we found that the use of HiFi-Cas9 for gene editing reduces the off-target INDEL and on-to-off-target translocation frequencies by more than 10-fold when compared to the WT-Cas9. Even though the compound treatment increases the off-target INDELs relative to the untreated sample, the frequency of off-target INDELs and translocations while using WT-Cas9 was substantially higher indicating that it is critical to use HiFi-Cas9 especially for therapeutic applications. Moreover, we observed that the increase in off-target INDEL frequency due to the compounds was dependent on their incubation times and reducing the incubation time to 12h post transfection was sufficient to normalize the INDELs to the levels of the untreated sample. This decreased incubation time also normalized the increase in the large on-target deletions observed with the 24 and 72h treatment. Thus, gene editing with HiFi Cas9 and a reduced incubation time of TFU72 can mitigate the potential genotoxic outcomes with minimal loss of HDR-enhancing activity. Another potential mitigation strategy for genotoxicity could be to combine the TFU72 treatment with an inhibitor of DNA polymerase theta which is a key player in the MMEJ repair pathway to reduce the large on-target deletions and translocations as described recently^31^. However, the safety of this approach to simultaneously inhibit NHEJ and MMEJ pathways is yet to be characterized through genome wide studies as it could have other unintended consequences.

Taken together, the novel DNA-PKcs inhibitor TFU72 is a potent and selective DNA-PKcs inhibitor that markedly enhances HDR gene editing across cell lines and therapeutically relevant human primary cells. Following the strategies we described for mitigating the potential genotoxic risks, TFU72 can be broadly applied for HDR-based targeted integration in the context of research and therapeutic applications.

## Methods

### Ethics statement

Umbilical cord blood-derived HSPCs were obtained from the Binns Program for Cord Blood Research with approval from the Stanford Institutional Review Board Committee under protocol 33813.

### Compounds and genome editing reagents

AZD7648 (Selleck Chemicals S8843, and MedChemExpress) was resuspended in DMSO to 2- or 10-mM stock solutions. Stock solution was stored at -20°C. The DNA-PK Inhibitor TFU72 was obtained from Merck Healthcare (Darmstadt, DE) as a 10 mM solution in DMSO. TFU72 will be available through Millipore Sigma as PURedit™ HDR Enhancer (HDRE01). For editing cell lines, PURedit Cas9 protein, synthetic single guide RNAs, and single stranded DNA donor templates (single-stranded oligodeoxynucleotides; ssODNs) all were purchased from MilliporeSigma (Burlington, MA). For editing primary cells, High-fidelity Cas9 was purchased from Aldevron (SpyFi Cas9, 9214) and WT Cas9 from IDT (Alt-R S.p. Cas9 Nuclease V3, 10000735). Chemically modified (2’-O-methyl-3’-phosphorothioate at the first and last three nucleotides) gRNAs were obtained from IDT and Synthego. Table S1 lists the genomic target sites of the gRNAs.

### Kinase panel testing

Biochemical kinase activity profiling was performed by Pharmaron (Beijing, China), according to the provider’s SOP using ADP-Glo (Promega) and HTRF KinEASE (Revity) formats in 384-well plates. TFU72 and AZD7648 were screened at 1 µM concentration across Pharmaron’s full kinase panel. Percent inhibition was calculated relative to plate vehicle and no-enyzme controls and runs met acceptance criteria (Z′ > 0.5; S/B > 2). Fluorescent signals were detected using EnVision 2104 or BMG multimode readers. A detailed protocol with all kinases and vendors is provided in Supplemental information.

### AAV6 vector construction, production and purification

The AAV transfer plasmid backbone pAAV-MCS (Agilent) was used for the construction of AAV transfer plasmids. Homology arms and donor sequences were cloned into the backbone using standard molecular biology techniques. AAV6 vectors were produced in-house or acquired through Vigene, Signagen, or Vectorbuilder. For in-house production, ∼10 million 293T cells were transfected with 22 μg packaging/helper plasmid pDGM6 (a gift from D. Russel, University of Washington, Addgene plasmid 110660) and 6 μg of transfer plasmid in 1 mL OptiMEM I (Gibco, 31985088) using PEI (Polysciences, 23966-1). After three days post-transfection, AAV6 was purified with the AAVpro purification kit (Takara, 6666). AAV6 titer was determined by ddPCR using a previously validated primer/probe set^45^ listed in Table S1.

### Culture and editing of cell lines (K562, HEK293 and U2OS)

K562, U2OS, and HEK293 cells were obtained from ATCC (Bethesda, MD) and grown at 37°C and 5% CO2 in their ATCC recommended media (MilliporeSigma) [Iscove’s Modified Dulbecco’s Medium for K562, McCoy’s 5a Medium Modified for U2OS, and Eagle’s Minimum Essential Medium for HEK293] supplemented with 10% FBS (MilliporeSigma) and GlutaMAX (ThermoFisher). Cells were seeded at 1.00 x 10^5^ cells/mL two days before transfection. At the time of transfection, cells were washed twice with Hank’s Balanced Salt Solution and resuspended in Nucleofector Solution for the corresponding cell type [SF for K562 and HEK293 and SE for U2OS] (Lonza, Walkersville, MD) at 10^6^ cells per 100 μL. Ribonucleoprotein (RNP) complexes were prepared by incubating for 15 minutes at room temperature: 30 pmol of PURedit Cas9 protein and 90 pmol of sgRNA in buffer (20 mM HEPES, 100 mM KCl, 0.5 mM DTT, 0.1 mM EDTA, pH 7.5). 300 pmol of the ssODN was then added for a total volume of 10 μL and then stored on ice until transfection. Nucleofection was performed by mixing 100 μL of prepared cell suspension with 10 μL of complexed RNP by pipetting up and down six times before transferring to a cuvette for electroporation using corresponding program on a Nucleofector 4d machine [FF-120 for K562, CM-104 for U2OS, and CM-130 for HEK293]. Nucleofected cells were immediately transferred to cellular media in a conical tube at 4.00 x10^5^ cells/mL. 1.00 x 10^5^ cells were then distributed to wells in 6-well plates containing media supplemented with the DNA-PK inhibitor to obtain wells containing 2000 μL with an inhibitor concentration of 1.0 µM. Some wells had no compounds for controls. Nucleofected cells were grown for 5 days before analysis and harvest. Guide Sequence, Donor Sequence, and NGS Primers are listed in Table S1.

### Cell viability analysis of cell lines (K562, HEK293 and U2OS)

Cell viability was tested with the CellTiter-Glo® 2.0 Assay (Promega, Madison, WI). An opaque-walled 96-well plate was prepared with 50 μL of each well from the 48-plate after thoroughly mixing and left at room temperature for 30 minutes. 50 μL of room temperature CellTiter-Glo® 2.0 Reagent was then added to each well and mixed on an orbital shaker for 10 minutes. Luminescence was then read on a SpectraMax iD3 (Danaher, Washington, DC).

### Gene editing quantification in cell lines (K562, HEK293 and U2OS) by NGS

Genomic DNA was harvested from the remaining cells using a MagMAX DNA Multi-Sample Ultra 2.0 Kit (ThermoFisher, Waltham, MA) following manufacturer protocol on a KingFisher Flex (96 deep-well format; ThermoFisher) or using the GenElute Mammalian Genomic DNA Miniprep Kit (Millipore, Burlington, MA). Genomic regions targeted by the RNP were amplified by PCR using NEBNext® Ultra™ II Q5® (NEB, Ipswich, MA) and the following cycling conditions: 98 C /30 s; 30 cycles of 98 C /10 s, 62 C /30 s, 65 C /45 s; 65 C /5 m. Primers are listed in Table 1. PCR products underwent a second round of amplification using Illumina index primers (Illumina, San Diego, CA) and NEBNext® Ultra™ II Q5® and the following conditions: 95 C /3m; 9 cycles of 95 C /30 s, 55 C /30 s, 72 C C/30 s; 72 C /5 m. Indexed PCR products were purified by Select-a-Size DNA Clean & Concentrator MagBeads (Zymo Research, Irvine, CA), using 1.2x beads by volume, quantified by PicoGreen (ThermoFisher), and pooled according to DNA content. Pools were diluted to 4 nM. Sequencing was performed on an Illumina MiSeq instrument (Illumina) using a 300-cycle kit to obtain single-end reads. FASTQ files for each sample were analyzed using Geneious software (GraphPad Software, LLC, Boston, MA), each file had at least 10,000 reads to be included in the data.

### iPSC culture and genome editing

Three human iPSC lines were utilized for gene editing experiments: PB005^46^, 1205-4^47^ and 1208-2^47^. PSCs were cultured feeder-free on Matrigel-coated plates (Corning, 354277) using mTeSR1 medium (STEMCELL Technologies, 85850). PSCs were pretreated for 24 h with 10 μM Y27632 (Cayman Chemicals, 10005583) prior to nucleofection. RNP complexes containing 6 μg Cas9 protein and 3.2 μg gRNA were complexed for 15 min at room temperature. Single-cell suspensions were generated using Accutase cell dissociation reagent (Innovative Cell Technologies, AT104). For nucleofection, 500,000 cells were resuspended in 20 μL P3 primary cell nucleofector solution (Lonza, V4XP-3032) containing the RNP complex, transferred to a 16-well Nucleocuvette Strip (Lonza), and nucleofected with program CA137 using a 4D-Nucleofector (Lonza). Following nucleofection, PSCs were seeded at 100,000 cells/well in 48-well plates using mTeSR1 containing 10 μM Y27632. AAV6 vectors and DNA-PKcs inhibitors were included in the culture medium at the specified concentrations. Fresh mTeSR1 with 10 μM Y27632 was used to replace the cell culture medium at 24 h post-nucleofection. One day after that, cells were cultured in mTeSR1 without any additives. Targeting efficiency was assessed between 4-6 days post-editing.

### CD34^+^ HSPC culture and genome editing

Human umbilical cord blood-derived CD34^+^ HSPCs were obtained through the Binns Program for Cord Blood Research. Cells were maintained in STEMSpan SFEM II (STEMCELL Technologies, 09655) containing 100 ng/mL stem cell factor (PeproTech, 300-07), 100 ng/mL thrombopoietin (PeproTech, 300-18), 100 ng/mL FLT3-ligand (PeproTech, 300-19), 100 ng/mL interleukin-6 (IL-6; Peprotech, 200-06), 20 U/mL penicillin, 20mg/mL streptomycin (Cytiva, SV30010) and 35 nM UM171 (APExBIO, A89505). Cells were maintained at 37 °C, 5% O_2_ and 5% CO_2_, with cell densities kept at 0.25-0.5 million cells/mL. RNP complexes for genome editing were generated by combining 6 μg of Cas9 with 3.2 μg of gRNA and allowing complex formation for 15 to 30 min at room temperature, then diluting into 20 μL of P3 primary cell nucleofector solution (Lonza, V4XP-3032). HSPCs (0.5-1 million) were resuspended in the 20 μL RNP-P3 mixture and nucleofected using program DZ-100 on a 4D-Nucleofector (Lonza). Nucleofected HSPCs were transferred to wells containing AAV6, AZD7648, and TFU72 at the specified concentrations. Fresh medium lacking AAV6 and small molecules was used to replace the culture medium after 24 h. Gene-targeting efficiency was evaluated at 2-3 days post-editing.

### CFU assay

For CFU assays, 500 HSPCs were plated 24 h post-editing in SmartDish six-well plates (STEMCELL Technologies, 27370) with containing MethoCult H4434 Classic (STEMCELL Technologies, 04444) according to the manufacturer’s instructions. After 14 d at 37 °C, 5% O_2_ and 5% CO_2_, colonies were counted and scored using a STEMvision automated counter (STEMCELL Technologies) to determine number of BFU-E, CFU-M, CFU-GM, and CFU-GEMM colonies.

### T cell culture and genome editing

Peripheral blood mononuclear cells (PBMCs) were isolated using Ficoll density gradient centrifugation, followed by negative selection of T cells with the EasySep™ Human T Cell Isolation Kit (StemCell Technologies, Vancouver, BC, Canada). Isolated T cells were cryopreserved in BamBanker media (GC Lymphotec, Tokyo, Japan) until further use.

Four days prior to electroporation, the cryopreserved T cells were thawed and cultured at a density of 1 x 10%6 cells/ml in a 24-well plate using T cell media: X-Vivo 15 (Lonza Bioscience, Basel, Switzerland), supplemented with 10% bovine growth serum (BGS) and recombinant human IL-2 (100 IU/ml, Peprotech/ThermoFisher, Waltham, MA, USA) at 37 °C, 5% CO_2_, and ambient O_2_. T cells were activated for three days with human T-Activator CD3/CD28 Dynabeads (Thermo Fisher) at a bead-to-cell ratio of 1:1. One day prior to electroporation, the beads were magnetically removed, and T cells were transferred to fresh media. Cell density was kept between 0.5-1 million cells/mL. Cells were washed with OptiMEM I prior to nucleofection. To prepare RNP complexes, 6 μg Cas9 was combined with and 3.2 μg gRNA and allowed to complex for 15-30 min at room temperature before diluting into 20 μL of P3 primary cell nucleofector solution (Lonza). Activated T cells (0.15-1 million) were resuspended in the RNP-P3 mixture and nucleofected using program EO-115 in 16-well strip cuvettes with a 4D-Nucleofector (Lonza). Following nucleofection, cells were placed in OptiMEM containing 100 IU/mL IL2 and AAV, AZD7648 and TFU72 at the indicated concentrations. Fresh XVIVO 15 medium (5% BGS, 100 IU/mL IL2) was provided after 24 h of culture. Editing efficiency was assessed at 2-3 days following the editing.

### Quantification of gene editing by ICE analysis

The ICE CRISPR analysis tool^48^ (EditCo Bio) was used to quantify the abundance and type of INDELs, and for short, targeted integrations the HDR frequency. In brief, gDNA was extracted from 50,000 to 200,000 cells using QuickExtract DNA Extraction Solution (QE; Lucigen, QE09050) and incubating at 65 °C for 6 min, followed by 100 °C for 10 min. 1-2 μL gDNA in QE were directly added to PCR reactions using either PrimeSTAR GXL DNA Polymerase (Takara, R050A), or Phusion Hot Start II High-Fidelity PCR Master Mix (Fisher Scientific, F566L), with primers that generate cut site-spanning amplicons. Amplified DNA was purified and Sanger-sequenced by MCLAB or GENEWIZ. The primer sequences are listed in Table S1.

### Quantification of allelic targeted integration by ddPCR analysis

The allelic frequence of target integration was measured by ddPCR analysis. For the ddPCR reaction, 2 μL of gDNA in QE were used in a 20 μL reaction containing 10 uL ddPCR supermix for probes (no dUTP, Biorad, 1863025), 0.5 μL of fluorophore-labeled target and reference primer/probe assays. The Bio-Rad droplet generator was used to generate ddPCR droplets with 20 μL ddPCR reaction mix and 70 μL droplet generation oil (Bio-rad, 1863005). The following PCR cycles were used with 40 μL of the generated droplet sample: 95 °C - 10 min, (94 °C - 30 s, 57 °C - 30 s, 72 °C - 2 min)*50, 98 °C – 10 min. The QX200 Droplet Digital PCR reader (BioRad) was used to analyze droplets after PCR. The number of positive target and reference droplets was determined using Quantasoft software (v1.7.4.0917). The allelic gene targeting frequency for each sample was calculated by dividing the number of target-positive droplets by the number of reference-positive droplets. The target primers were designed as described previously^2^, with an In-Out PCR strategy where one primer anneals to the integrated sequence and the other outside of the homology arm. The primer and probe sequences are listed in Table S1.

### Quantification of cellular targeted integration by flow cytometry

For edits that involved targeted knock-in of large GFP-encoding constructs, flow cytometry was performed to measure the percentage of GFP+ cells 3 days post-editing. iPSCs were analyzed using a BD FACSAria II Flow Cytometer; HSPCs and T-cells were measured with a BD LSRFortessa Flow Cytometer.

### Cell viability analysis for iPSCs and HSPCs

Cell viability of gene-edited PSCs was determined using an MTT assay. Following gene editing, PSCs were seeded into 96-well plates and viability was measured at 24, 48, and 72 h post-editing. MTT (MedChemExpress, HY-15924) was diluted to 0.5 mg/mL in growth medium and added to cells and incubated for 2 h at 37 °C. Following incubation, the MTT solution was aspirated, and cells were lysed with 100 μl lysis buffer containing 0.1 N HCl and 0.5% SDS in isopropanol. Absorbance measurements were taken at 570 nm (reference wavelength 650 nm) using a SpectraMax M3 plate reader (Molecular Devices). Cell viability was expressed as a percentage relative to mock-treated controls based on the absorbance readings. HSPC cell viability was determined through viable cell count using TC10 counter (Biorad) following resuspension in equal volume of Trypan blue (Gibco, 15250061).

### Genotoxicity Assessment

Off-target INDELs at HBB-OT1 site in chromosome 9 was performed through NGS analysis. For this, we used a commercial sequencing service Amplicon-EZ through Azenta (Genewiz). The sequence spanning of the off-target site was PCR amplified with illumina adapter added to the primers. The PCR amplicon was purified using Genejet PCR purification kit. The amplicon was submitted to Azenta at a concentration of 20 ng μl–1, in 25-μl volume. DNA library preparation and sequencing was performed at Azenta. DNA library was prepared using the NEBNext Ultra DNA Library Prep kit following the manufacturer’s instructions. Limited cycle PCR was used to index and enrich the DNA amplicons. DNA libraries validated on Agilent TapeStation (Agilent Technologies) and multiplexed in equal molar mass following quantification using a Qubit 2.0 fluorometer (Invitrogen). Pooled DNA libraries were sequenced using a 2 × 250-bp paired-end configuration using an Illumina sequencer. Raw FastQ files from the NGS was analyzed by using the CRISPResso2 tool for quantification of percent reads with INDELs following the instructions from the developer.

To quantify the chromosomal translocation between the HBB on target and off-target site 1, we designed ddPCR primer/probe to assess four different translocation outcomes. The translocation frequency was calculated as the sum of the frequency of all 4 events normalized to a reference locus (LCR) in the same chromosome as the HBB locus (Chr11). The primers/probe used for translocation ddPCR are listed in the Table S1.

For large deletion analysis, we used the KOD -Multi & Epi-™ high-fidelity DNA polymerase^49^ (Toyobo) to amplify a 10-kb sequence spanning the HBB on-target site using the primer pair listed in Table S1. This polymerase has been shown to be ideal for large deletion analysis as it has minimal size bias during PCR amplification. The amplicons were purified using the Genejet PCR purification kit and was submitted to Plasmidsaurus for the standard premium PCR oxford nanopore sequencing. The raw FASTQ files from the sequencing was analyzed as described in a previous publication^50^. Since no code was provided in the original publication, generative AI was used to reimplement the described analysis pipeline from scratch. Accuracy of the code was validated by using synthetic data with exactly defined deletion size distribution profiles.

### Statistical Analysis

Statistical analyses were performed using Graphpad Prism10 software. Two-way ANOVA with multiple comparisons test was used for the analysis.

## Generative AI use and declaration

During the preparation of this work the author(s) used Claude (Anthropic) and ChatGPT (OpenAI) in order to help refine text and to write the code for large deletion analysis. After using this tool/service, the author(s) reviewed and edited the content as needed and take(s) full responsibility for the content of the published article.

## Data Availability

NGS dataset will be made available at the time of publication along with the code used for analysis of large deletions.

## Acknowledgements

We thank the Binns Program for Cord Blood Research for providing cord blood CD34+ HSPCs. We thank the FACS Core Facility at the Stanford Institute of Stem Cell Biology and Regenerative Medicine for access to the FACS machines. M.H.P. acknowledges funding from Sutardja Chuk Professorship and Laurie Lacob Endowment. L.S. acknowledges funding from the Knut and Alice Wallenberg Foundation. F.K.E. acknowledges Stanford Knight-Hennessy Scholarship, Paul and Daisy Soros Fellowship for New Americans, and the Hertz Fellowship.

