## Supplemental information for "TFU72 is a novel and potent DNA-PKcs inhibitor for enhancing homology-directed repair gene editing"

### WT full panel (384 targets) Kinase Activity Assay protocol

#### 1 Material and Instrument

**Table 1 Experimental Materials Information**

| Materials and Reagents | Vendor |
|---|---|
| ADP-Glo™ Kinase Assay | Promega |
| HTRF KinEASE-TK kit | Revvity |
| HTRF KinEASE -STK S1 kit | Revvity |
| HTRF KinEASE -STK S2 kit | Revvity |
| HTRF KinEASE -STK S3 kit | Revvity |
| DMSO | Sigma |
| DTT (DL-Dithiothreitol) | Sigma |
| $MgCl_2$ | Sigma |
| HEPES | Solarbio |
| BRIJ 35 detergent (10%) | Millipore |
| EGTA Solution | Macklin |
| BSA stabilizer 7.5% | Revvity |
| $MnCl_2$ | Sigma |
| NaCl | Sigma |
| Trition X-100 (10%) | Thermo Scientific |
| $CaCl_2$ | Sigma |
| $MnCl_2$ | Sigma |
| CDK2/CyclinO | Biortus |
| CDK2/CyclinD1 | Biortus |
| CDK1/CyclinE2 | Biortus |
| LTK | Carna |
| MNK1 | Carna |

| | |
|---|---|
| MNK2 | Carna |
| SIK1 | Carna |
| SIK2 | Carna |
| SIK3 | Signalchem |
| DMPK1 | Carna |
| LOK | Carna |
| MST1 | Carna |
| GPRK4 | Carna |
| ULK3 | Carna |
| ALK6 | Eurofins |
| CK1 alpha 1 | Signalchem |
| CK1 delta | Signalchem |
| CK1 epsilon | Signalchem |
| CK1 gamma 2 | Signalchem |
| CK1 gamma 3 | Signalchem |
| PLK1 | Carna |
| PLK2 | Carna |
| PLK3 | Signalchem |
| PLK4 | Signalchem |
| CK1 alpha 1L | Signalchem |
| CK2 alpha 1 | Signalchem |
| CK2 alpha 2 | Signalchem |
| GRK5 | Biortus |
| mTOR | Invitrogen |
| TLK1 | Carna |
| TLK2 | Carna |
| VRK2 | Eurofins |
| ALK2 | Signalchem |

| | |
|---|---|
| ALK4 | Carna |
| CK1 gamma 1 | Signalchem |
| CK2a1/b | Carna |
| CK2a2/b | Carna |
| IKK epsilon | Signalchem |
| NEK7 | Carna |
| PKMYT1 | Carna |
| CDK6/CyclinD2 | Biortus |
| BRSK1 | Carna |
| BRSK2 | Carna |
| MARK1 | Carna |
| MARK2 | Carna |
| MARK3 | Carna |
| MARK4 | Carna |
| NIM1K | Carna |
| NUAK1 | Signalchem |
| NUAK2 | Carna |
| TSSK1 | Carna |
| PKN1/PRK1 | Signalchem |
| PKN2/PRK2 | Signalchem |
| TNK1 | Signalchem |
| PIK3CA/PIK3R1 | Millipore |
| CDK4/CyclinD1 | Invitrogen |
| DYRK4 | Carna |
| DYRK1A | Carna |
| DYRK1B | Signalchem |
| DYRK2 | Carna |
| DYRK3 | Carna |

| | |
|---|---|
| SRPK2 | Carna |
| MST4 | Carna |
| MSSK1 | Carna |
| MEK1 | Signalchem |
| MEK2 | Signalchem |
| CDK3/CyclinE2 | Signalchem |
| PKD2 | Carna |
| PKD3 | Carna |
| MST3 | Carna |
| VEGFR3 | Carna |
| TAK1 | Carna |
| BARK2 | Carna |
| GSK3 alpha | Carna |
| GSK3 beta | Signalchem |
| CDK1/CyclinA2 | Signalchem |
| CDK1/CyclinB1 | Proqinase |
| CDK1/CyclinE1 | Biortus |
| GPRK7 | Carna |
| GPRK6 | Signalchem |
| CDk2/CyclinE1 | Carna |
| CDK2/CyclinE2 | Biortus |
| CDK2/CyclinA1 | Signalchem |
| CDK2/CyclinA2 | Invitrogen |
| CDk3/CyclinE1 | Signalchem |
| CDK4/CyclinD3 | Carna |
| CDK5/p25 | Signalchem |
| CDk5/p35 | Signalchem |
| CDK6/CyclinD1 | ThermoFisher |

| | |
|---|---|
| CDK6/CyclinD3 | Carna |
| GAK | Biortus |
| SLK | Signalchem |
| MAPKAPK3 | Carna |
| MAPKAPK5 | Carna |
| MK2 | Signalchem |
| Jak2 (JH1, JH2 domain) | BPS |
| IKK beta | Signalchem |
| JAK1 | Invitrogen |
| TYK2 | Invitrogen |
| LRRK2 | ThermoFisher |
| BMPR2 | Carna |
| CAMKK2 | Signalchem |
| CDK18/CyclinY | Biortus |
| CDK7/CycH/MAT1 | Biortus |
| CLK1 | Carna |
| CLK2 | Signalchem |
| CLK4 | Carna |
| CRIK | Signalchem |
| Erk1 | Signalchem |
| Erk2 | Signalchem |
| ERK5 | Carna |
| ERK7 | Carna |
| ERN1 | Signalchem |
| GCK | Carna |
| GLK | Invitrogen |
| HGK | Carna |
| HIPK1 | Signalchem |

| | |
|---|---|
| HIPK2 | Carna |
| HIPK3 | Signalchem |
| HIPK4 | Signalchem |
| HPK1 | Signalchem |
| ICK | Carna |
| KHS | Signalchem |
| LIMK1 | Carna |
| MAP3K14 | Carna |
| MEKK2 | Carna |
| MEKK3 | Carna |
| MINK1 | Signalchem |
| TNIK | Signalchem |
| MLK1 | Carna |
| MLK2 | Signalchem |
| MLK3 | Invitrogen |
| MST2 | Carna |
| NEK1 | Carna |
| NEK2 | Carna |
| NEK4 | Carna |
| NEK9 | Carna |
| NLK | Carna |
| PKR | Carna |
| CDK1/CyclinA1 | Signalchem |
| AMPK alpha1/beta1/gamma1 | Carna |
| CaMK2 gamma | Carna |
| CaMK2 delta | Carna |
| CAMK2 beta | Carna |
| MYLK2 | Signalchem |

| | |
|---|---|
| CaMK1 alpha | Carna |
| CaMK1 delta | Carna |
| CaMK2 alpha | Carna |
| DCAMKL2 | Carna |
| DAPK1 | Carna |
| DAPK3 | Carna |
| CHEK1 | Signalchem |
| CHEK2 | Signalchem |
| DRAK1 | Carna |
| PRKCA | Carna |
| RSK1 | Signalchem |
| RSK2 | Signalchem |
| RSK3 | Signalchem |
| IRAK4 | Carna |
| Aurora B | Carna |
| Aurora A | Signalchem |
| Aurora C | Carna |
| PAK4 | Carna |
| PAK6 | Biortus |
| PAK7 | Carna |
| PKAC alpha | Signalchem |
| PKAC beta | Signalchem |
| PKAC gamma | Signalchem |
| RSK4 | Carna |
| PRKX | Carna |
| AKT1 | Signalchem |
| AKT2 | Signalchem |
| AKT3 | Signalchem |

| | |
|---|---|
| PIM2 | Carna |
| SGK1 | Signalchem |
| PIM3 | Signalchem |
| SGK2 | Carna |
| p70S6K | Signalchem |
| PIM1 | Carna |
| PKD1 | Carna |
| ROCKI | ThermoFisher |
| ROCKII | ThermoFisher |
| PKG1A | Millipore |
| TAOK1 | ThermoFisher |
| TAOK2 | Carna |
| TBK1 | Carna |
| TGFBR2 | Biortus |
| TSSK2 | Carna |
| TTBK1 | Carna |
| TTBK2 | Carna |
| TTK | Invitrogen |
| ULK1 | Signalchem |
| ULK2 | Signalchem |
| WNK1 | Carna |
| WNK2 | Carna |
| WNK3 | Carna |
| YSK4 | Signalchem |
| TPL2 | Carna |
| CDK4/CyclinD2 | Biortus |
| AAK1 | Signalchem |
| IRAK1 | Carna |

| | |
|---|---|
| NDR1 | Carna |
| NDR2 | Carna |
| p38 alpha | Signalchem |
| p38 gamma | Carna |
| p38 beta | Signalchem |
| p38 delta | Signalchem |
| JNK1 | Carna |
| JNK2 | Carna |
| JNK3 | Carna |
| MAP2K3 | Carna |
| MAP2K6 | Carna |
| PAK1 | Carna |
| CDC7/DBF4 | Signalchem |
| CDK9/CycK | Biortus |
| CDK9/CycT1 | Biortus |
| CDK9/CycT2 | Signalchem |
| PDK1 | Signalchem |
| PIP5K1A | Signalchem |
| TAOK3 | Carna |
| GRK2 | Signalchem |
| FRK | Signalchem |
| SRM | Carna |
| ACK | Signalchem |
| JAK3 | Invitrogen |
| TEC | Carna |
| TXK | Signalchem |
| ASK1 | Biortus |
| Wee1 | BPS |

| | |
|---|---|
| CDK12/CycK | Biortus |
| CDK13/CycK | Biortus |
| OSR1 | Carna |
| BUB1/BUB3 | Carna |
| RIPK3 | Vkeybio |
| MSK1 | Carna |
| MSK2 | Carna |
| DMPK2 | Carna |
| MRCK alpha | Eurofins |
| MRCK beta | Eurofins |
| LATS1 | Carna |
| LATS2 | Carna |
| PAK2 | Carna |
| PAK3 | Carna |
| SGK3 | Carna |
| SPHK1 | Signalchem |
| SPHK2 | Signalchem |
| ALK5 | Signalchem |
| cRAF (Raf1) | Carna |
| MELK | Signalchem |
| PASK | Carna |
| PHKG1 | Carna |
| PHKG2 | Carna |
| MKK7 | Carna |
| TAK1-TAB1 | Signalchem |
| AMPK alpha1/beta1/gamma2 | Signalchem |
| AMPK alpha1/beta1/gamma3 | Signalchem |
| AMPK alpha1/beta2/gamma2 | Signalchem |

| | |
|---|---|
| AMPK alpha2/beta1/gamma2 | Signalchem |
| AMPK alpha2/beta1/gamma3 | Signalchem |
| AMPK alpha2/beta2/gamma2 | Signalchem |
| ROS1 | Signalchem |
| TIE2 | Signalchem |
| MUSK | Carna |
| VEGFR2 | Signalchem |
| EPHA1 | Carna |
| EPHA2 | Signalchem |
| EPHA3 | Carna |
| EPHA5 | Carna |
| EPHA7 | Carna |
| EPHA8 | Carna |
| EPHB2 | Carna |
| EPHB3 | Carna |
| EPHB4 | Carna |
| PDGFR alpha | Carna |
| PDGFR beta | ThermoFisher |
| ABL2 | Signalchem |
| AXL | Signalchem |
| BMX | Signalchem |
| LYN A | Signalchem |
| LYN B | Signalchem |
| FGR | Signalchem |
| ABL1 | Signalchem |
| FAK | Signalchem |
| LCK | Signalchem |
| FES | Signalchem |

| | |
|---|---|
| TYRO3 | Carna |
| EPHA4 | Carna |
| EPHB1 | Carna |
| DDR2 | Signalchem |
| DDR1 | ThermoFisher |
| PYK2 | Signalchem |
| PKC beta1 | Carna |
| PKC beta2 | Carna |
| PKC gamma | Carna |
| PKC delta | Carna |
| PKC epsilon | Carna |
| PKC zeta | Carna |
| PKC eta | Carna |
| PKC theta | Carna |
| PKC iota | Carna |
| DCAMKL1 | Carna |
| RIPK1 | Biortus |
| RIPK2 | Signalchem |
| SOK1 | Signalchem |
| BRK | Carna |
| Her4 | Signalchem |
| GCN2 | Signalchem |
| SYK | Signalchem |
| TRKB | Invitrogen |
| VEGFR1 | Invitrogen |
| ZAP70 | Carna |
| IGF1R | Signalchem |
| INSR | Signalchem |

| | |
|---|---|
| EGFR | Signalchem |
| Met | Carna |
| FLT3 | Carna |
| c-Kit | Carna |
| ALK | Carna |
| RON | Signalchem |
| CSF1R | Biortus |
| CSK | Carna |
| IRR | Carna |
| ITK | Signalchem |
| FYN A | Signalchem |
| HCK | Signalchem |
| RET | Carna |
| SRC | Signalchem |
| YES | Signalchem |
| FGFR1 | Invitrogen |
| FGFR2 | Invitrogen |
| FGFR3 | Invitrogen |
| FGFR4 | ThermoFisher |
| TRKC | Invitrogen |
| TRKA | Signalchem |
| BLK | Signalchem |
| c-MER | ThermoFisher |
| FYN B | Carna |
| Her2 | ThermoFisher |
| EPHA6 | Carna |
| PIP5K1C | Signalchem |
| PIP5KL1 | Carna |

| | |
|---|---|
| PI4K2A | Signalchem |
| PI4K2B | Signalchem |
| PI4KB | Signalchem |
| PIP5K3 | Carna |
| PIK3CG | Invitrogen |
| DGKA | Signalchem |
| DGKB | Signalchem |
| DGKE | Signalchem |
| DGKQ | Signalchem |
| AMPK alpha2/beta1/gamma1 | Carna |
| IKK alpha | Biortus |
| FER | Carna |
| MAP3K4 | Carna |
| CLIK1 | Biortus |
| DLK | Carna |
| PI42A | Biortus |
| CDK19/CyclinC/MED12 | Biortus |
| ACTR2 | Signalchem |
| ACTR2B | Signalchem |
| LKB1 | Carna |
| VRK1 | Biortus |
| PKG1 | Biortus |
| PKG2 | Biortus |
| SRPK1 | Biortus |
| YANK1 | Biortus |
| TSSK4 | Biortus |
| Wee1B | Biortus |
| SBK | Biortus |

| | |
|---|---|
| BIKE | Biortus |
| JAK2 | Invitrogen |
| ZAK | Carna |
| BRAF | Carna |
| BTK | Signalchem |
| CLK3 | Carna |
| AMPK alpha1/beta2/gamma3 | Signalchem |
| PIK3C2A | Invitrogen |
| PIK3C2G | Invitrogen |
| HRI | Carna |
| DGKZ | Signalchem |
| ATM | Eurofins |
| ATR | Eurofins |
| DNA-PK | Eurofins |
| PIP4K2B | Carna |

**Table 2 Instruments and Consumables Information**

| Instruments and Consumables | Vendor |
|---|---|
| 384 polystyrene shallow flat white | Greiner |
| 384-Well Polypropylene microplate | Labcyte |
| Plate shaker | Thermo |
| Centrifuge | Cence |
| High-throughput drug screening kinase reader | BMG |
| Envision 2104 multilabel Reader | PerkinElmer |
| Echo 655 | Labcyte |

### 2 Experimental Methods

#### 2.1 Steps of Kinase Experiments by the ADP-Glo Method

a) Preparation of 1× kinase Buffer.

b) Transfer compound dilutions into each well of assay plates (784075, Greiner) using Echo 655.

c) Seal the assay plate, centrifuge compound plates at 1000 rpm for 1 min.

d) Prepare 2× Enzyme in 1× kinase buffer.

e) Add 2.5μL of 2× Enzyme into 384-well assay plate except the positive control (784075, Greiner).

f) Add 2.5 μL of 1× kinase buffer into the wells of positive control (784075, Greiner).

g) Centrifuge plates at 1000 rpm for 1 min, RT for 10 min.

h) Prepare 2× Substrate and ATP mixture in 1× kinase buffer.

i) Start the reaction by adding 2.5 μL 2× Substrate and ATP mixture.

j) Centrifuge plates at 1000 rpm for 1 min. Seal the assay plate, RT for 2 h.

k) Add 4 μL ADP-Glo reagents. Incubate at RT for 60 min.

l) Add 8 μL kinase detection reagents. Incubate at RT for 60 min. m) Read Luminescence signal on Envision 2104 or BMG plate reader.

#### 2.2 Steps of Kinase Experiments by the TR-FRET Method

a) Preparation of 1× kinase Buffer.

b) Transfer compound dilutions into each well of assay plates (784075, Greiner) using Echo 655.

c) Seal the assay plate, centrifuge compound plates at 1000 rpm for 1 min.

d) Prepare 2× Enzyme in 1× kinase buffer.

e) Add 5 μL of 2× Enzyme into 384-well assay plate except the positive control (784075, Greiner).

f) Add 5 μL of 1× kinase buffer into the wells of positive control (784075, Greiner).

g) Centrifuge plates at 1000 rpm for 1 min, RT for 10 min.

h) Prepare 2× TK-substrate-biotin or 2× STK-substrate-biotin and ATP mixture in 1× kinase buffer.

i) Start the reaction by adding 5 μL TK-substrate-biotin or STK-substrate-biotin and ATP.

j) Centrifuge plates at 1000 rpm for 1 min. Seal the assay plate, RT for 40 min. k) Add 5 μL Sa-XL 665 and 5μL TK-antibody-Cryptate or 5 μL STK-antibody-Cryptate mixture to the corresponding wells of the assay plate.

l) Centrifuge plate at 1000 rpm for 1 min, RT for 1 h.

m) Read fluorescence signal at 615 nm (Cryptate) and 665 nm (XL665) on Envision 2104 or BMG plate reader.

**2.3 Data analysis**

a) Inhibition Percentage Calculation

%Inhibition =100-(Signal cmpd – Signal $_{Ave\_PC}$) / (Signal $_{Ave\_VC}$-Signal $_{Ave\_PC}$) ×100

Signal $_{Ave\_PC}$: the average value of the positive control (no kinase added) on the reaction plate.

Signal $_{Ave\_VC}$: the average value of the negative control (kinase reaction solution without test article) on the reaction plate.

b) Assay acceptance criteria

Z factor > 0.5; S/B > 2.

S/B = mean signal value of the negative control / mean signal value of the positive control

Z factor =1-3* (Stdev value of the negative control + Stdev value of the positive control) / (mean of the negative control - mean of the positive controls)

### Figure S1

#### A Kinase panel: inhibition (1 μM) at 20 to 50% (sorted for TFU72)

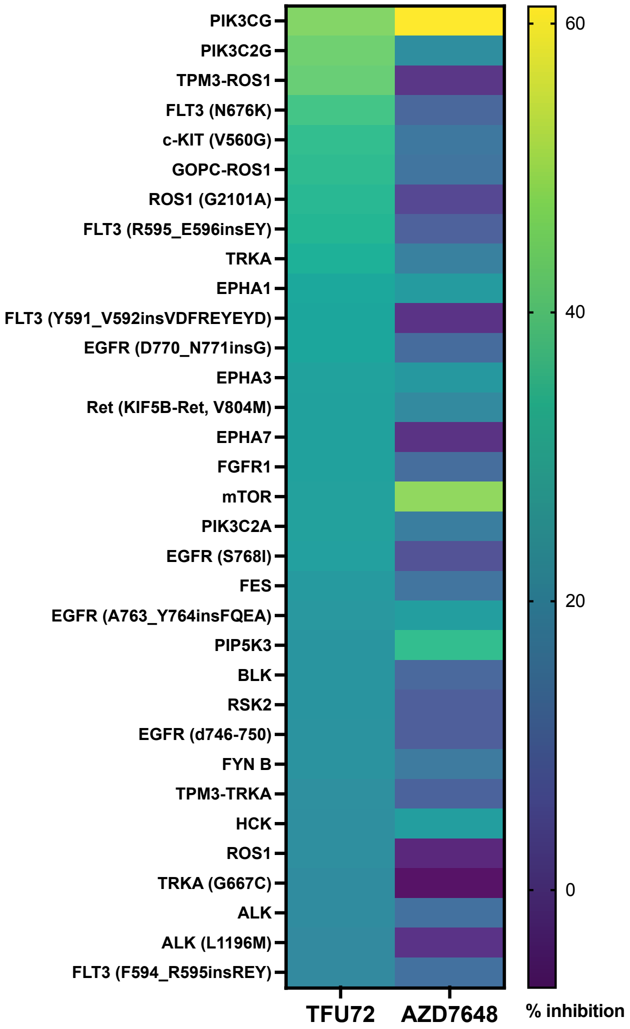


#### B Kinase panel: inhibition (1 μM) at 20 to 50% (sorted for AZD7648)

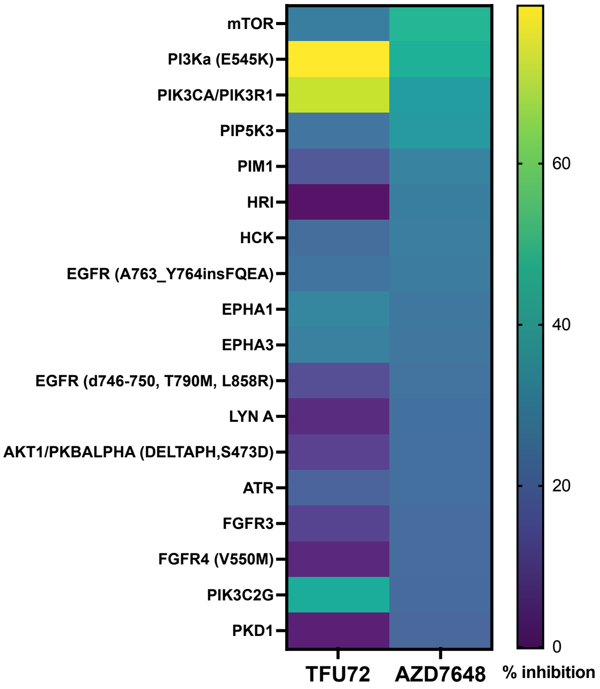

**Figure S1. Kinases partially inhibited by TFU72 and AZD7648. A-B.** Heatmap showing the top kinases inhibited by TFU72 and AZD7648 (both at 1 μM) following testing on a human kinase panel. The data is shown in descending order sorted for kinases inhibited at 20-50% relative to a target-specific inhibitor by TFU72 (**A**) or AZD7648 (**B**). The kinases are sorted in descending order for % incubation by TFU72 (A) or AZD7648 (B).

### Figure S2

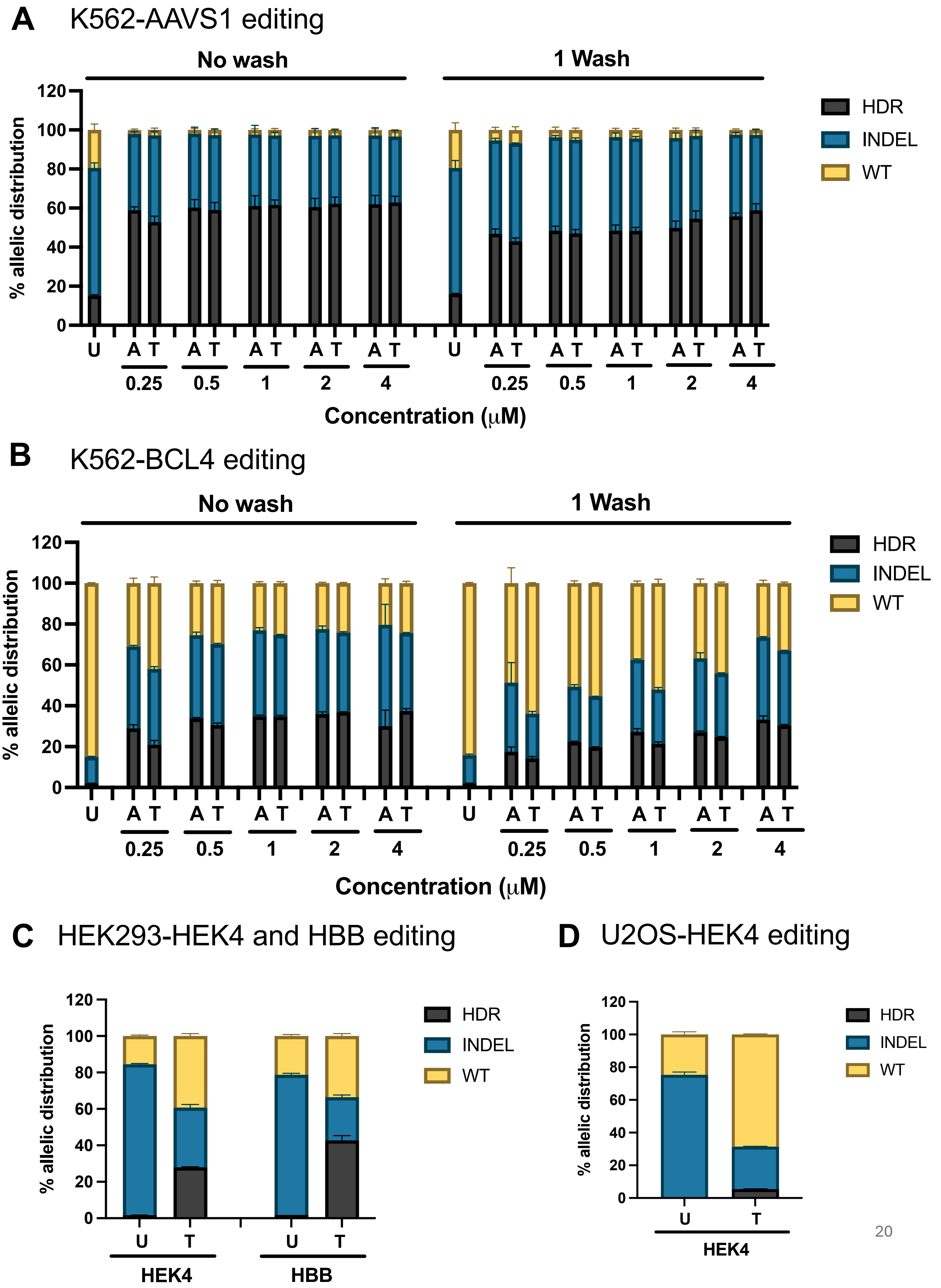
A K562-AAVS1 editing
No wash
1 Wash
% allelic distribution
HDR
INDEL
WT
U A T A T A T A T A T U A T A T A T A T A T
0.25 0.5 1 2 4 0.25 0.5 1 2 4
Concentration (μM)
B K562-BCL4 editing
No wash
1 Wash
% allelic distribution
HDR
INDEL
WT
U A T A T A T A T A T U A T A T A T A T A T
0.25 0.5 1 2 4 0.25 0.5 1 2 4
Concentration (μM)
C HEK293-HEK4 and HBB editing
% allelic distribution
HDR
INDEL
WT
U T U T
HEK4 HBB
D U2OS-HEK4 editing
% allelic distribution
HDR
INDEL
WT
U T
HEK4

**Figure S2. TFU72 improves HDR editing across genomic loci and in various cell lines. A-B.** Allelic distribution of HDR, INDEL and WT frequencies in K562 cells edited at AAVS1 (**A**) and BCL4 (**B**) loci using RNP and ssODN with or without AZD7648 and TFU72 treatment as assessed by NGS. U denotes untreated (editing without compound), A and T denote editing with AZD7648 and TFU72 (1 μM), respectively (n=2). The compounds were removed either 24 hours post editing (1 wash) or left in the cells for 5 days (0 wash). **C-D**. Allelic distribution of HDR, INDEL and WT frequencies in HEK293 edited at HEK4 and HBB loci (**C**) and U2OS cells edited at HEK4 locus (**D**) using RNP and ssODN with or without TFU72 treatment as assessed by NGS. U denotes untreated (editing without compound) and T denote editing with TFU72 (1 μM) (n=2).

### Figure S3

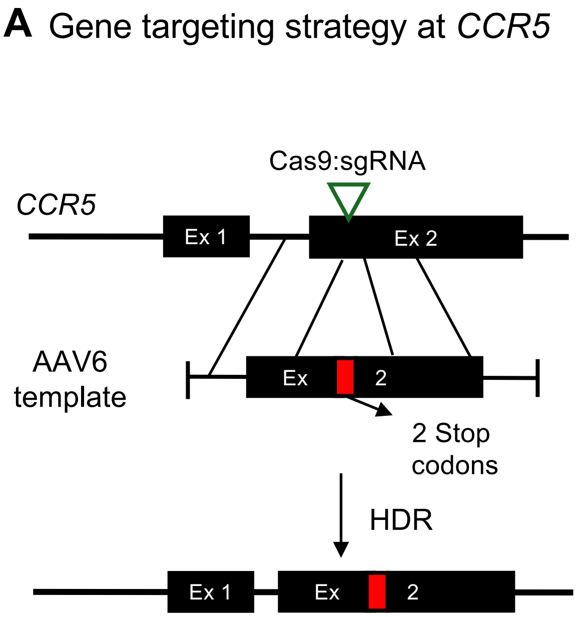
A Gene targeting strategy at CCR5
Cas9:sgRNA
CCR5
Ex 1
Ex 2
AAV6 template
Ex
2
2 Stop codons
HDR
Ex 1
Ex
2

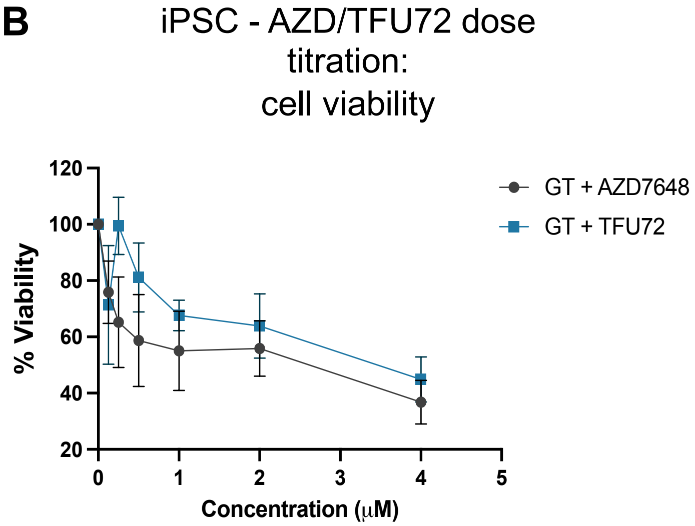
B
iPSC - AZD/TFU72 dose titration: cell viability
GT + AZD7648
GT + TFU72
% Viability
Concentration (μM)

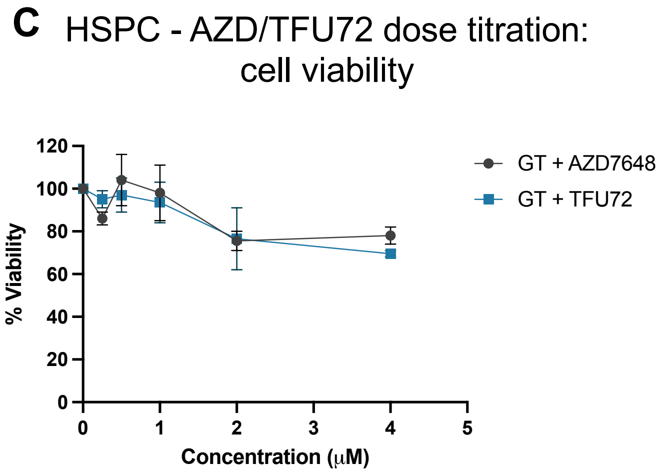
C
HSPC - AZD/TFU72 dose titration: cell viability
GT + AZD7648
GT + TFU72
% Viability
Concentration (μM)

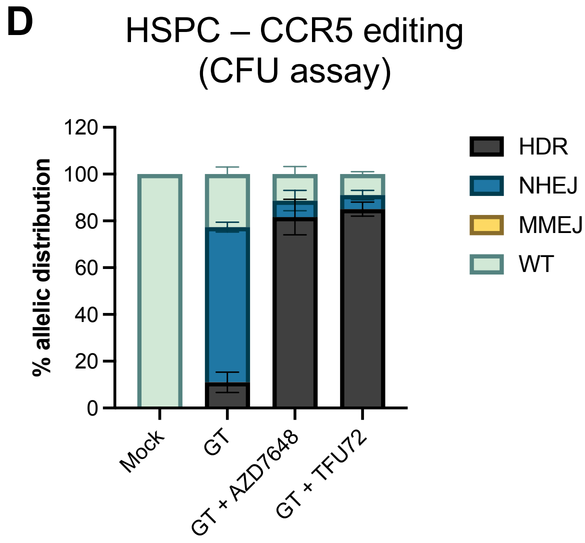
D
HSPC – CCR5 editing (CFU assay)
HDR
NHEJ
MMEJ
WT
% allelic distribution
Mock
GT
GT + AZD7648
GT + TFU72

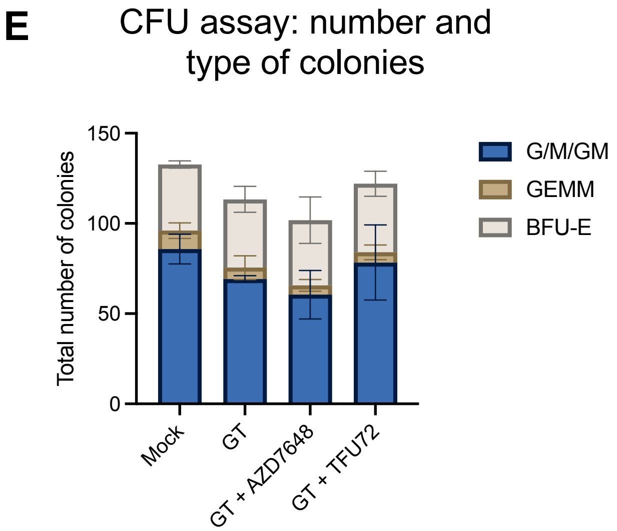
E
CFU assay: number and type of colonies
G/M/GM
GEMM
BFU-E
Total number of colonies
Mock
GT
GT + AZD7648
GT + TFU72

**Figure S3. Viability of iPSC and HSPC edited with TFU72. A.** Schematic showing gene targeting strategy at the CCR5 locus for the knock-in of two stop codons. **B**. Cell viability of iPSC gene edited at CCR5 locus with different concentrations of AZD7648 and TFU72 was assessed by MTT cell viability assay. The data is plotted as viability relative to the cells edited without the compounds. GT denotes gene targeting (HDR) with RNP, AAV6 (n=2). The HDR frequency of the corresponding samples is shown in Figure 2A-B. **C**. Cell viability of HSPC gene edited at CCR5 locus with different concentrations of AZD7648 and TFU72 was assessed by measuring viable cell count at D3 post editing. The data is plotted as viability relative to the cells edited without the compounds (n=2). **D-E.** HSPCs were gene edited at CCR5 locus for the knock-in of short sequence (two stop codons) using RNP and AAV6 gene editing with or without AZD7648/TFU72 compounds (1 μM). Bar graph shows allelic distribution of HDR, NHEJ, MMEJ and WT frequencies (**D**) (n=3). Colony forming units (CFU) assay was performed on the CCR5 gene edited HSCs to assess the viability and differentiation potential. Data is shown as the distribution of different colony types and the corresponding number of colonies **(E).** GEMM denotes granulocyte, erythroid, macrophage, megakaryocyte, GM denotes granulocyte and monocyte, and BFU-E denotes burst forming unit-erythroid (n=3).

### Figure S4

**A** Gene targeting strategy at *HBB*

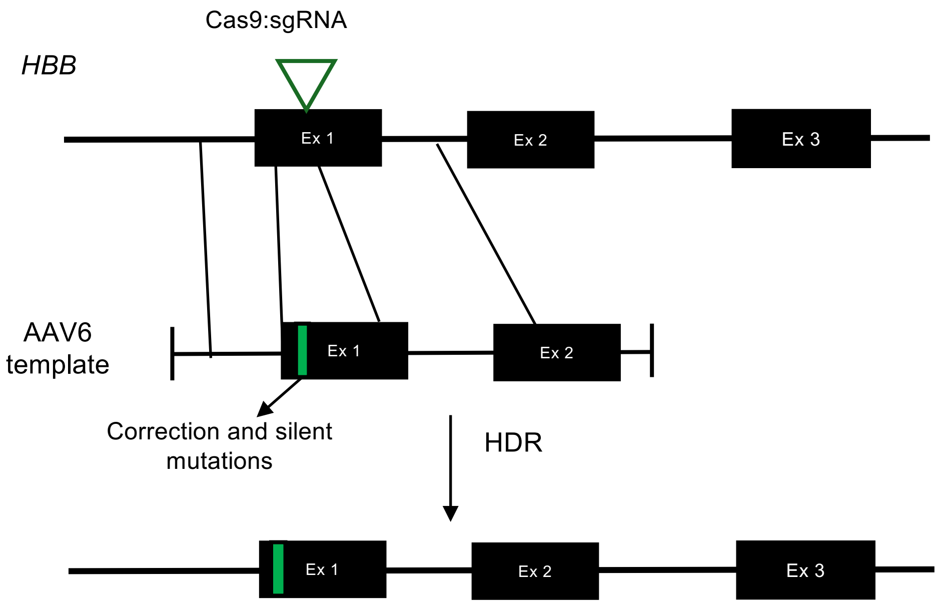


**B** Gene targeting strategy at *STING1*

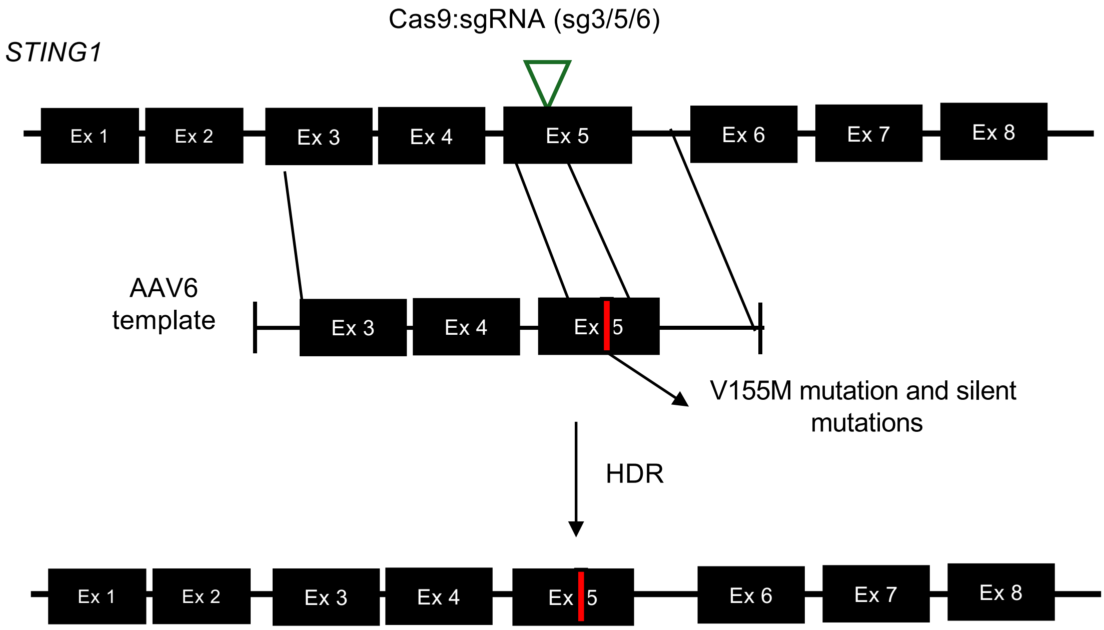

### Figure S4

C iPSC - short insert KI – NHEJ, MMEJ and WT frequencies

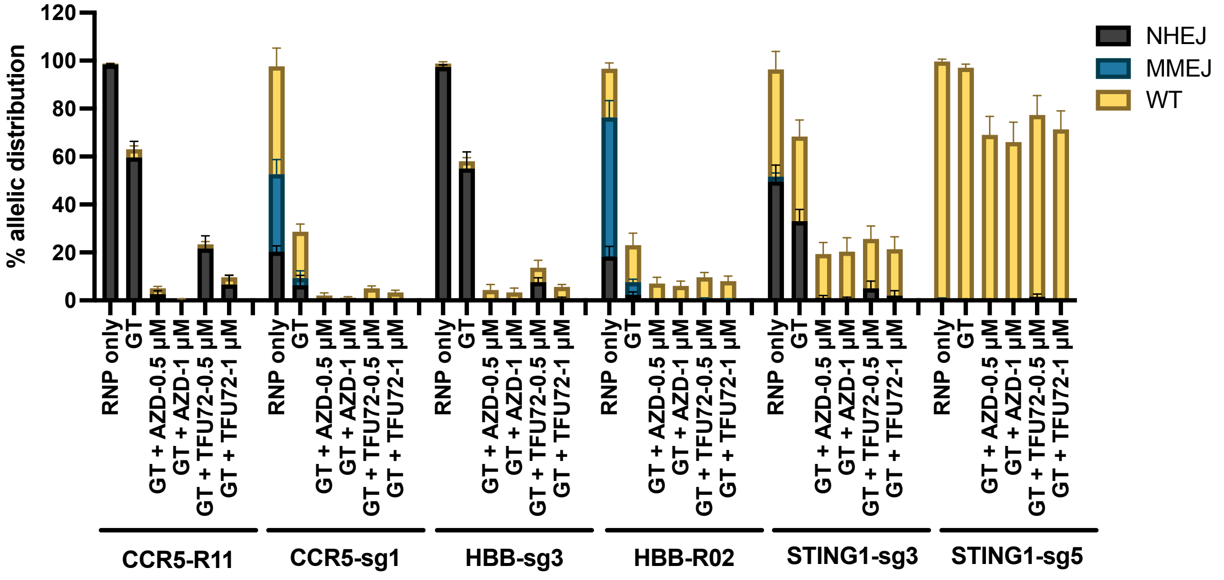


D HSPC - short insert KI – NHEJ, MMEJ and WT frequencies

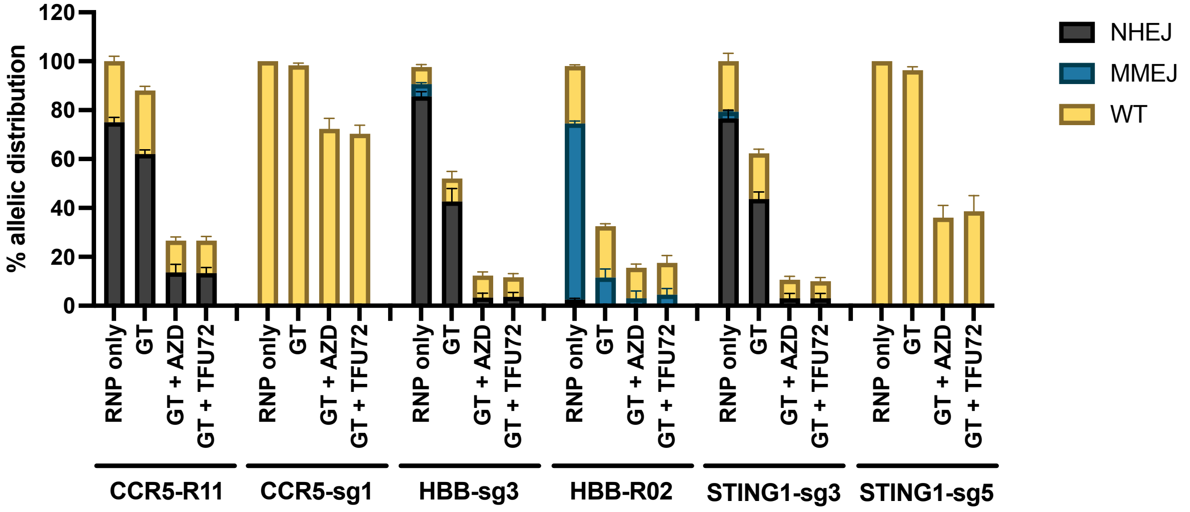


E T cells - short insert KI – NHEJ, MMEJ and WT frequencies

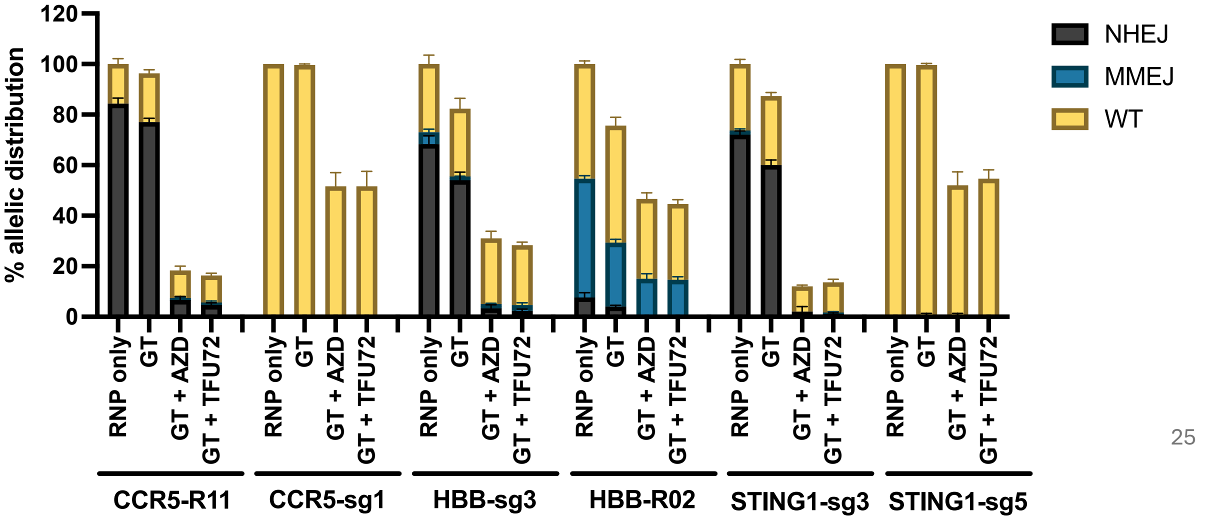

**Figure S4. WT and INDEL frequency in iPSC, HSPC and T cells HDR edited with TFU72. A-B.** Schematic for gene targeting strategy at HBB **(A)** and STING1 **(B)** loci for the correction of sickle cell disease mutation (HBB) and introduction of V155M point mutation (STING1), respectively. **C-E.** iPSC (**C**), HSPC (**D**) and T cells (**E**) were individually edited at CCR5, HBB and STING1 loci with two different gRNAs at each loci using RNP and AAV6 gene editing for the knock-in of short sequence with AZD7648 and TFU72 at the indicated concentrations. Bar graphs show distribution of allelic NHEJ INDEL, MMEJ INDEL and WT frequencies measured at D3 post editing using Sanger Sequencing and ICE analysis (n=3). HDR frequencies of the corresponding samples is shown in Figure 3. GT denotes gene targeting (HDR) with RNP, AAV6 editing. AZD denotes AZD7648. The concentration of AZD7648 and TFU72 used for iPSC were 0.5 μM and 1 μM as indicated while for HSPC and T cells, the compounds were used at 0.5 and 1 μM, respectively. AAV6 HDR template for CCR5 locus was designed to knock-in two stop codons at the target site. At the HBB locus, the AAV6 HDR template was designed to knock-in silent mutations and correction of the sickle cell disease mutation (E6V). For the STING1 locus, the AAV6 HDR template was designed to knock-in a point mutation (V155M) along with a few silent mutations at the target site.

### Figure S5

**A** Large insert knock-in strategy at *CCR5*

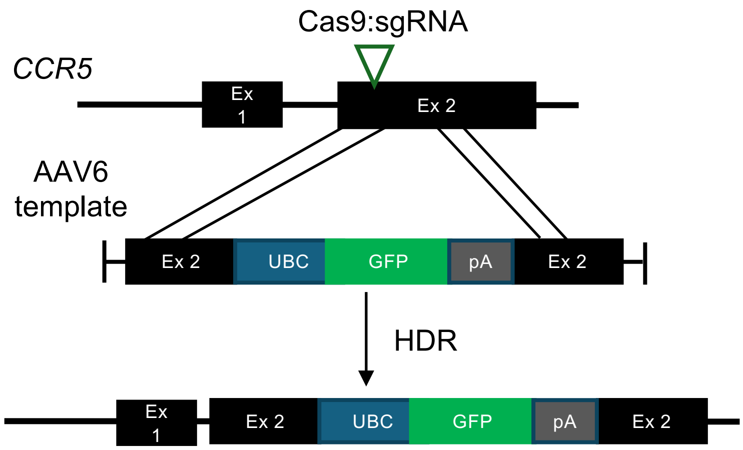


**B** Large insert knock-in strategy at *HBB*

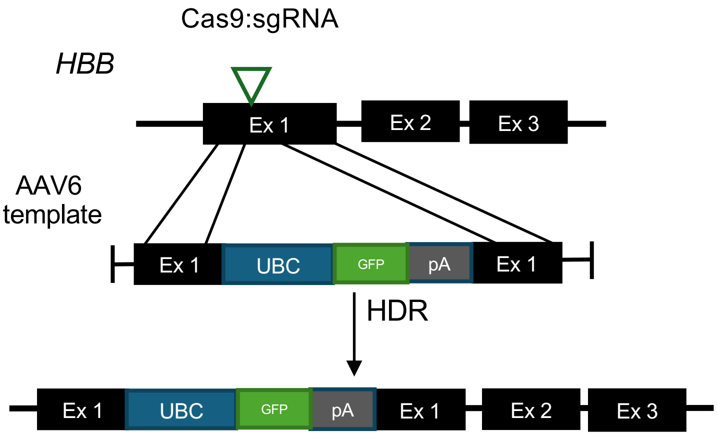


**C** Large insert knock-in strategy at *STING1*

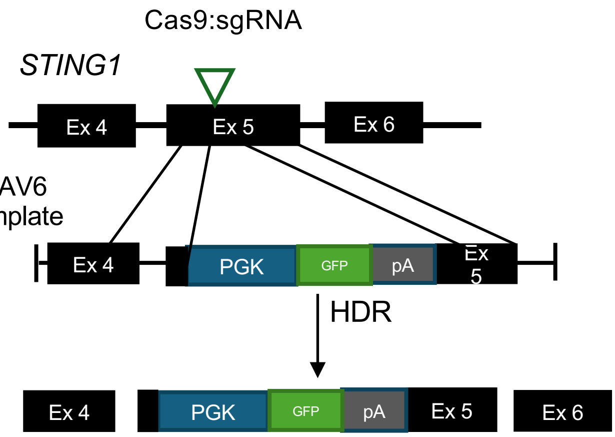

**Figure S5. Schematics for large insert knock-in strategy. A-C.** Schematics showing the strategy for large insert knock-in for GFP expression at *CCR5* (**A**)**,** *HBB* (**B**) and *STING1* (**C**) loci using RNP/AAV6 gene editing.

### Figure S6

**A** iPSC - large insert KI and AAV6 MOI titration: flow cytometry

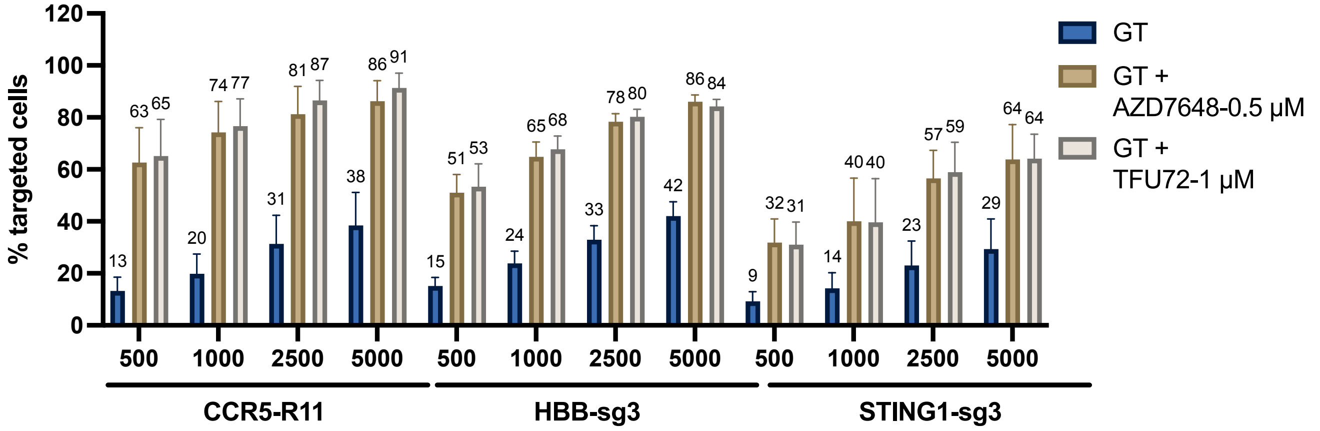


**B** iPSC - large insert KI and AAV6 MOI titration: WT and INDEL type distribution

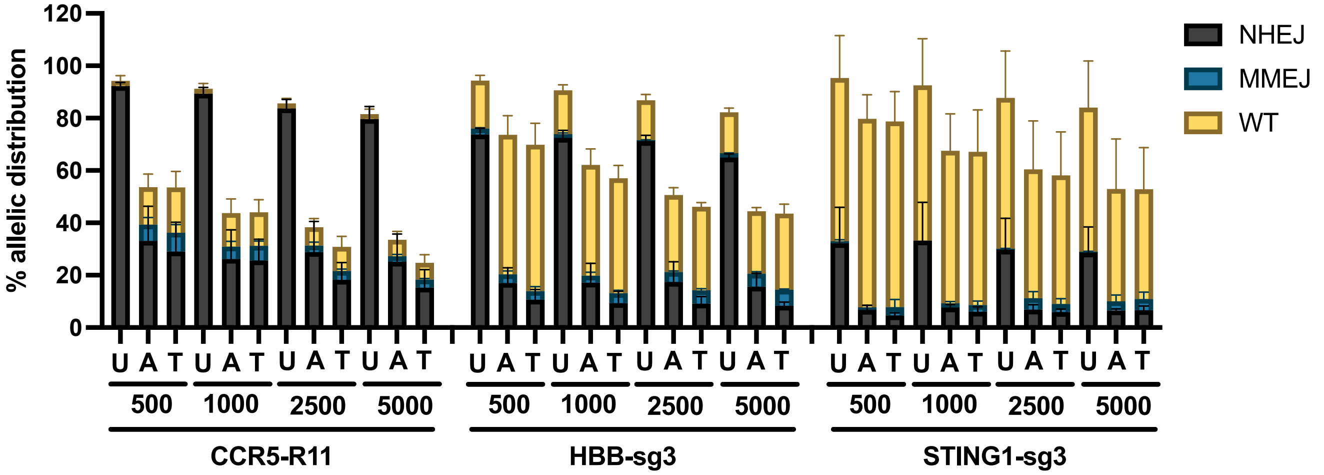


**C** HSPC - large insert KI and AAV6 MOI titration: flow cytometry

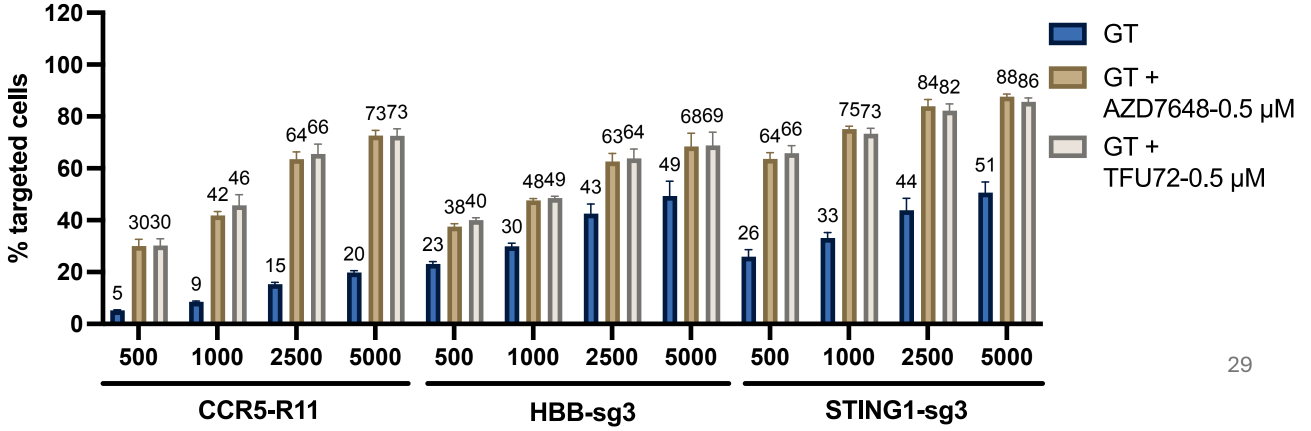

### Figure S6

**D** HSPC - large insert KI and AAV6 MOI titration: WT and INDEL type distribution

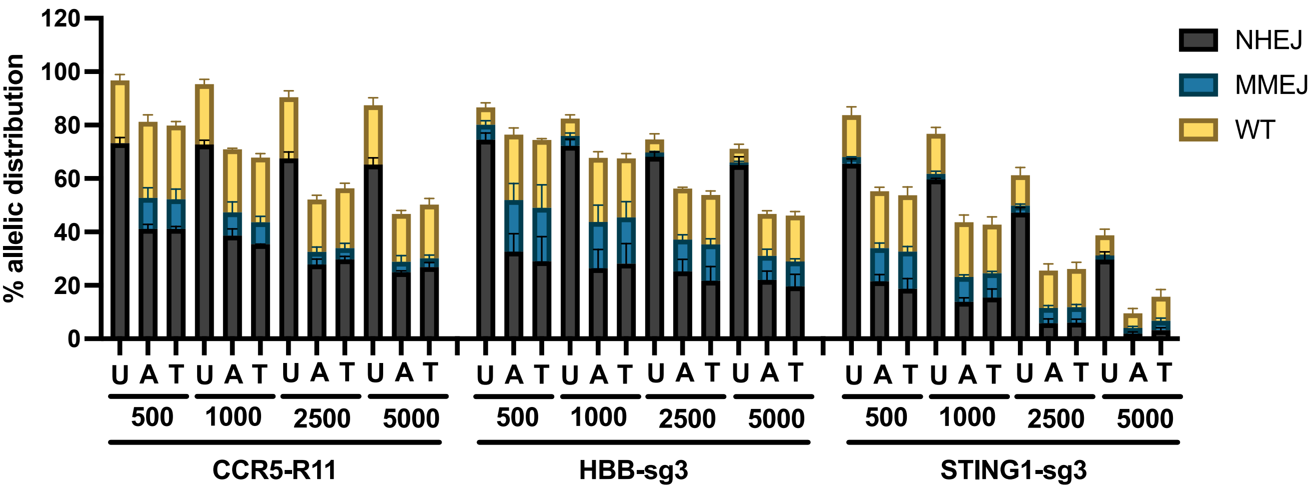


**E** T cells - large insert KI and AAV6 MOI titration: flow cytometry

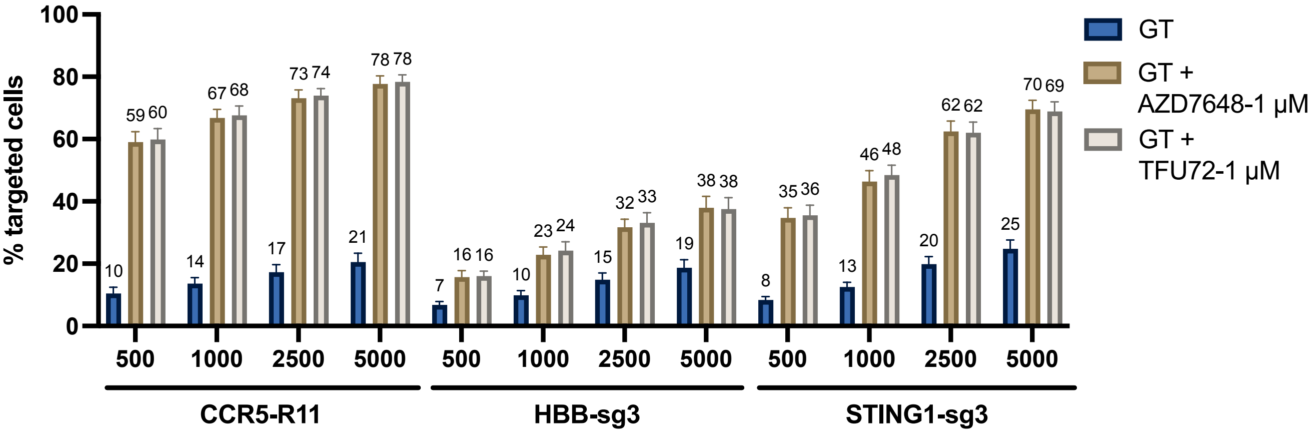


**F** T cells - large insert KI and AAV6 MOI titration: WT and INDEL type distribution

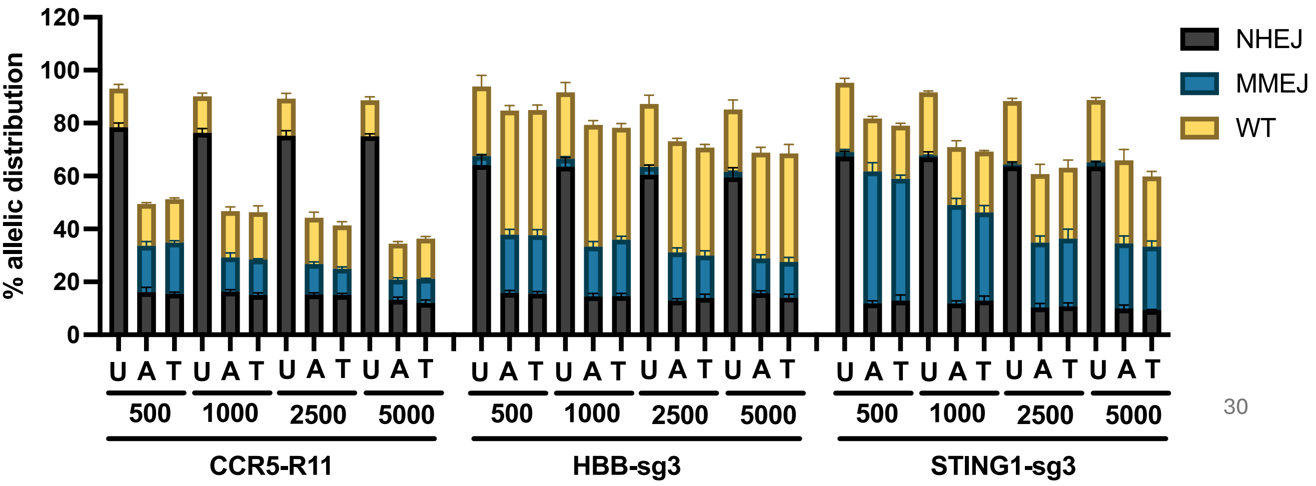

**Figure S6. HDR editing with TFU72 for large insert KI with AAV6 MOI titration in iPSC, HSPC and T cells. A-F.** iPSC (**A, B**), HSPC (**C, D**) and T cells (**E, F**) were edited at CCR5, HBB and STING1 loci individually using RNP and AAV6 gene editing for the knock-in of multi-kb sequence with AZD7648 and TFU72 at the indicated concentrations and at different AAV6 multiplicity of infection (MOI): 500, 1000, 2500, 5000. **A, C, E.** Bar graphs show frequency of HDR edited cells (% targeted cells) as measured at D3 post editing by flow cytometry for GFP (n=3). **B, D, F.** Bar graphs show distribution of allelic NHEJ INDEL, MMEJ INDEL and WT frequencies in the HDR edited cells (n=3). The corresponding allelic HDR frequencies are shown in Figure 4. GT denotes gene targeting (HDR) with RNP, AAV6 editing. AAV6 HDR templates for CCR5 and HBB loci were designed to knock-in a 2.2-kb sequence consisting of UBC promoter driven GFP followed by a bGH polyA signal sequence. For the STING1 locus, the AAV6 HDR template was designed to knock-in a 1.4-kb sequence consisting of PGK promoter driven GFP followed by a sv40 polyA signal sequence.

#### Figure S7

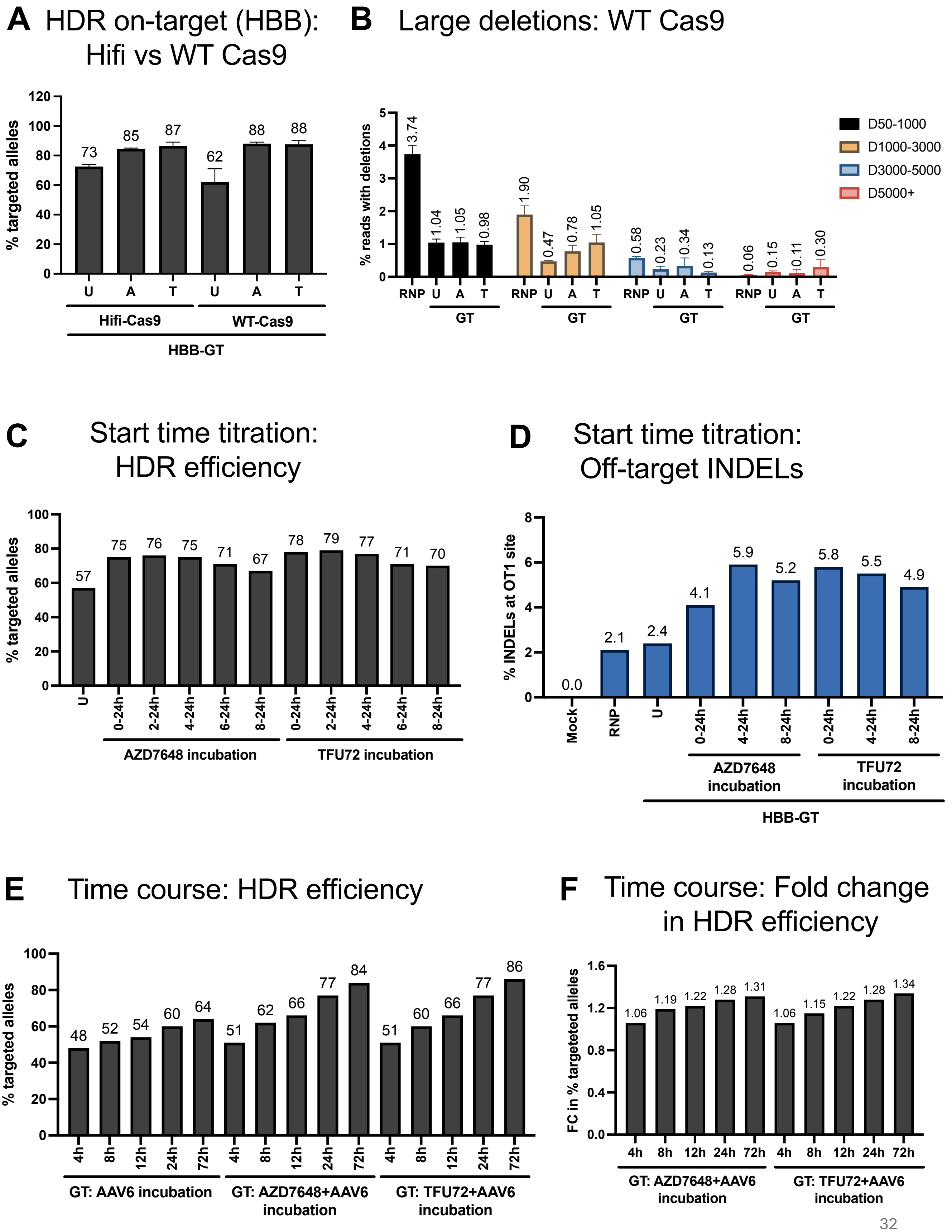

A HDR on-target (HBB): Hifi vs WT Cas9
% targeted alleles
73 85 87 62 88 88
U A T U A T
Hifi-Cas9
WT-Cas9
HBB-GT
B Large deletions: WT Cas9
% reads with deletions
D50-1000
D1000-3000
D3000-5000
D5000+
3.74 1.04 1.05 0.98 1.90 0.47 0.78 1.05 0.58 0.23 0.34 0.13 0.06 0.15 0.11 0.30
RNP U A T
GT
C Start time titration: HDR efficiency
% targeted alleles
57 75 76 75 71 67 78 79 77 71 70
U 0-24h 2-24h 4-24h 6-24h 8-24h 0-24h 2-24h 4-24h 6-24h 8-24h
AZD7648 incubation
TFU72 incubation
D Start time titration: Off-target INDELs
% INDELs at OT1 site
0.0 2.1 2.4 4.1 5.9 5.2 5.8 5.5 4.9
Mock RNP U 0-24h 4-24h 8-24h 0-24h 4-24h 8-24h
AZD7648 incubation
TFU72 incubation
HBB-GT
E Time course: HDR efficiency
% targeted alleles
48 52 54 60 64 51 62 66 77 84 51 60 66 77 86
4h 8h 12h 24h 72h
GT: AAV6 incubation
GT: AZD7648+AAV6 incubation
GT: TFU72+AAV6 incubation
F Time course: Fold change in HDR efficiency
FC in % targeteted alleles
1.06 1.19 1.22 1.28 1.31 1.06 1.15 1.22 1.28 1.34
4h 8h 12h 24h 72h
GT: AZD7648+AAV6 incubation
GT: TFU72+AAV6 incubation

**Figure S7. Characterization of gene editing with TFU72 and incubation time optimization.** HSPCs were edited at HBB locus for the HDR editing of SCD mutation using RNP, AAV6 (Hifi or WT Cas9) with AZD7648 (A, 0.5 μM) or TFU72 (T, 0.5 μM) or no treatment (U). GT denotes gene targeting (HDR) with RNP, AAV6. **A**. Bar graph shows HDR frequency at HBB locus as determined by ICE analysis (n=2). Corresponding off-target INDELs, on- to off-target translocation and Hifi Cas9 large deletion data is shown in Figure 5A-C. **B**. HSPCs edited at HBB loci with WT Cas9 were assessed for the frequency of large deletions at the on-target site using Nanopore sequencing of a 10-kb PCR amplicon. The data is shown as bar graph of frequency of reads with deletions of the sizes 50-1000, 1000-3000, 3000-5000 and above 5000 bp (n=2). **C-D.** HSPCs edited at HBB locus with AZD7648 and TFU72 at different incubation start times post editing were assessed for on-target HDR by ICE analysis and off-target INDELs were analyzed by NGS at D3 post editing. **C.** Bar graph shows the frequency of targeted alleles (HDR). **D.** Bar graph shows the frequency of INDELs at the OT1 off-target site as determined by the CRISPResso2 tool analysis of the NGS data. **E-F.** HSPCs edited at HBB locus with different incubation times of AAV6, AAV6+AZD7648 and AAV6+TFU72 were assessed for HDR frequency at D3 post editing by ICE analysis. **E.** Bar graph shows the frequency of allelic HDR efficiency (% targeted alleles). **F.** Bar graph shows the fold change in the frequency of HDR alleles in AZD7648 or TFU72 treated samples relative to the corresponding untreated samples with same AAV6 incubation times.
